# Urgency reveals an attentional vortex during antisaccade performance

**DOI:** 10.1101/433615

**Authors:** Emilio Salinas, Benjamin R Steinberg, Lauren A Sussman, Sophia M Fry, Christopher K Hauser, Denise D Anderson, Terrence R Stanford

**Affiliations:** Department of Neurobiology and Anatomy, Wake Forest School of Medicine, 1 Medical Center Blvd., Winston-Salem, NC 27157-1010, USA

**Keywords:** attention, capture, decision making, individual differences, mental chronometry, salience

## Abstract

In the antisaccade task, which is considered a sensitive assay of cognitive function, a salient visual cue appears and the participant must look away from it. This requires sensory, motor-planning, and cognitive neural mechanisms. But what are the unique contributions of these mechanisms to performance, and when exactly are they engaged? By introducing an urgency requirement into the antisaccade task, we track the evolution of the choice process with millisecond resolution and find a singular, nonlinear dependence on cue exposure: when viewed briefly (∼100–140 ms), the cue captures attention so powerfully that looking at it (erroneously) is virtually inevitable, but as the cue viewing time increases, the probability of success quickly rises and saturates. The psychophysical and modeling results reveal concerted interactions between reflexive and voluntary cognitive mechanisms that (1) unfold extremely rapidly, (2) are qualitatively consistent across participants, and (3) are nevertheless quantitatively distinctive of each individual’s perceptual capacities

---

Neuroscience aims to explain macroscopic behavior based on the microscopic operation of distinct neural circuits, and this requires carefully designed tasks that expose the relationship between the two. In the case of the antisaccade task (Coe and Munoz, 2017; Munoz and Everling, 2004), in which participants are instructed to withhold responding to a salient visual stimulus in favor of programming a saccade to a diametrically opposed location, performance relies heavily on frontal cortical mechanisms associated with cognitive control (Guitton et al., 1985; Everling and Fischer, 1998; Munoz and Everling, 2004; Condy et al., 2007; Luna et al., 2008; Hakvoort et al., 2012). The paradigm is considered to be a sensitive assay of impulsivity and executive function in general, and indeed, the mean reaction time (**RT**) and overall error rate in the antisaccade task are frequently used as biomarkers for cognitive development (Klein and Foerster, 2001; Luna et al., 2008; Coe and Munoz, 2017) and, in clinical settings, for mental dysfunction (Everling and Fischer, 1998; Munoz et al., 2003; Hutton and Ettinger, 2006; Antoniades et al., 2015; Wiecki et al., 2016).

The antisaccade task pits against each other two fundamental processes, one involuntary and the other voluntary. On one hand, the sudden appearance of the cue automatically attracts spatial attention (Theeuwes, 1991; Theeuwes et al., 1998; Ruz and Lupiáñez, 2002; Theeuwes, 2010; Carrasco, 2011; Aagten-Murphy and Bays, 2017). The attraction can result in a covert shift (“attentional capture”) or an overt saccade (“oculomotor capture”), but in either case the effect is described as bottom-up or exogenous, and is thought to be fast and transient. On the other hand, programming a saccade away from the cue summons top-down or endogenous attention, and perhaps other mechanisms (e.g., working memory [Roberts et al., 1994; Lavie and De Fockert, 2005]). This process is thought to be slower and to require a sustained cognitive effort (Godijn and Theeuwes, 2002; Theeuwes, 2010; Carrasco, 2011). Thus, the rationale for the task is sound — the timing and intensity of the conflict between bottom-up and top-down mechanisms should correlate with behavior, and with the dynamics of the underlying attentional and oculomotor neural circuits.

There is a problem, however: such conflict must unfold very quickly. First, exogenous attention is thought to be mediated by visually-driven responses in oculomotor areas such as the frontal eye field (**FEF**) and superior colliculus (**SC**), which have latencies of at least 50 ms (Gottlieb and Goldberg, 1999; Thompson et al., 2005; Ipata et al., 2006; Joiner et al., 2017; White et al., 2017; Chen et al., 2018). And second, spatial attention can be endogenously shifted roughly 150 ms after a relevant cue is provided (Kim and Cave, 1999; Ogawa and Komatsu, 2004; Busse et al., 2008; Theeuwes, 2010; Markowitz et al., 2011). This leaves less than 100 ms for the competition between exogenous and endogenous responses to evolve. The usual behavioral metrics of mean RT and overall accuracy are thus unlikely to yield a clean characterization of this competition, because they constitute the average end results of numerous operations (perceptual, motor, cognitive) that contribute to a much longer choice process (indeed, below we show that such metrics are severely confounded). How can this problem be overcome?

**Figure 1.**
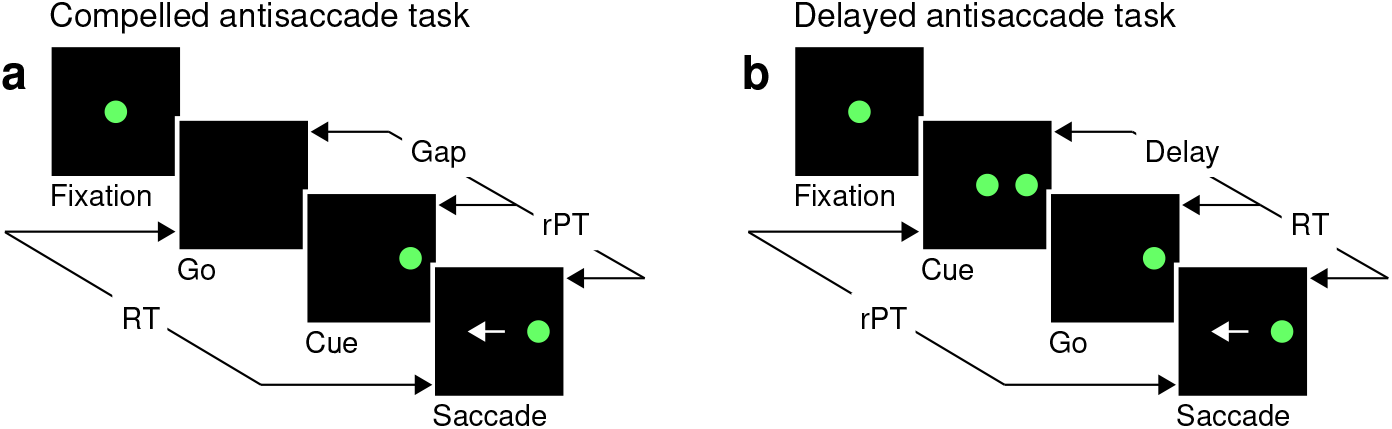
Urgent and non-urgent variants of the antisaccade task. **a**, The compelled antisaccade task. After a fixation period (150, 250, or 350 ms), the central fixation point disappears (Go), instructing the participant to make an eye movement to the left or to the right (±10^°^) within 450 ms. The cue is revealed (Cue) after a time gap that varies unpredictably across trials (Gap, 0–350 ms). The correct response is an eye movement (Saccade, white arrow) away from the cue, to the diametrically opposite, or anti, location. **b**, The delayed antisaccade task. In this case the cue is shown before the go signal, during fixation. The interval between cue onset and fixation offset varies across trials (Delay, 100 or 200 ms). In all trials, the reaction time (RT) is measured between the onset of the go signal and the onset of the saccade, whereas the raw processing time (rPT) is measured between cue onset and saccade onset.

The solution is to make the task urgent. The compelled antisaccade task requires subjects, humans in this case, to begin programming a saccade before knowing the direction of the correct response, and to use later arriving information about cue location to appropriately modify the ongoing motor plans. Urgency allows us to generate a special psychometric function, the “tachometric curve,” which tracks success rate as a function of the perceptually relevant time interval, the raw processing time (**rPT**, measured between cue presentation and saccade onset). We find that, for the compelled antisaccade task, the tachometric curve takes on a unique shape: within a narrow rPT range, the curve yields a pronounced dip to below-chance performance — an attentional vortex — in which the capture by the cue is so strong that the success rate approaches 0%; thereafter, however, the fraction of correct saccades to the ‘anti’ location increases extremely rapidly. The experimental data were comprehensively replicated by a neurophysiologically-based model of the saccadic choice process, with the combined results providing a remarkably detailed account of how reflexive and voluntary mechanisms compete over time to determine task performance.

## Results

### Urgent antisaccade behavior is characterized by an attentional vortex

Akin to an athlete anticipating the trajectory of a ball that must be caught or struck, the participant in the compelled antisaccade task must begin programming a movement in advance of the relevant sensory information, and must quickly interpret the later arriving visual cue to modify the developing motor plan(s) on the fly. In the sequence of task events (Fig. 1a), the key step is the early offset of the fixation spot, which means “respond now!” This go signal is given first, before the cue, which is revealed after an unpredictable gap period. The cue appears randomly to the left or to the right of fixation, and the participant is instructed to make an eye movement away from it, to the diametrically opposite location — but for this response to be correct, the saccade must be initiated within 450 ms of the go signal. The urgent nature of our compelled antisaccade task, in which motor and perceptual processes are meant to run concurrently, stands in contrast to the easy, non-urgent version of the task (Fig. 1b), in which delivery of the cue before the go signal allows more time for the perceptual process to be completed before saccade onset.

**Figure 2.**
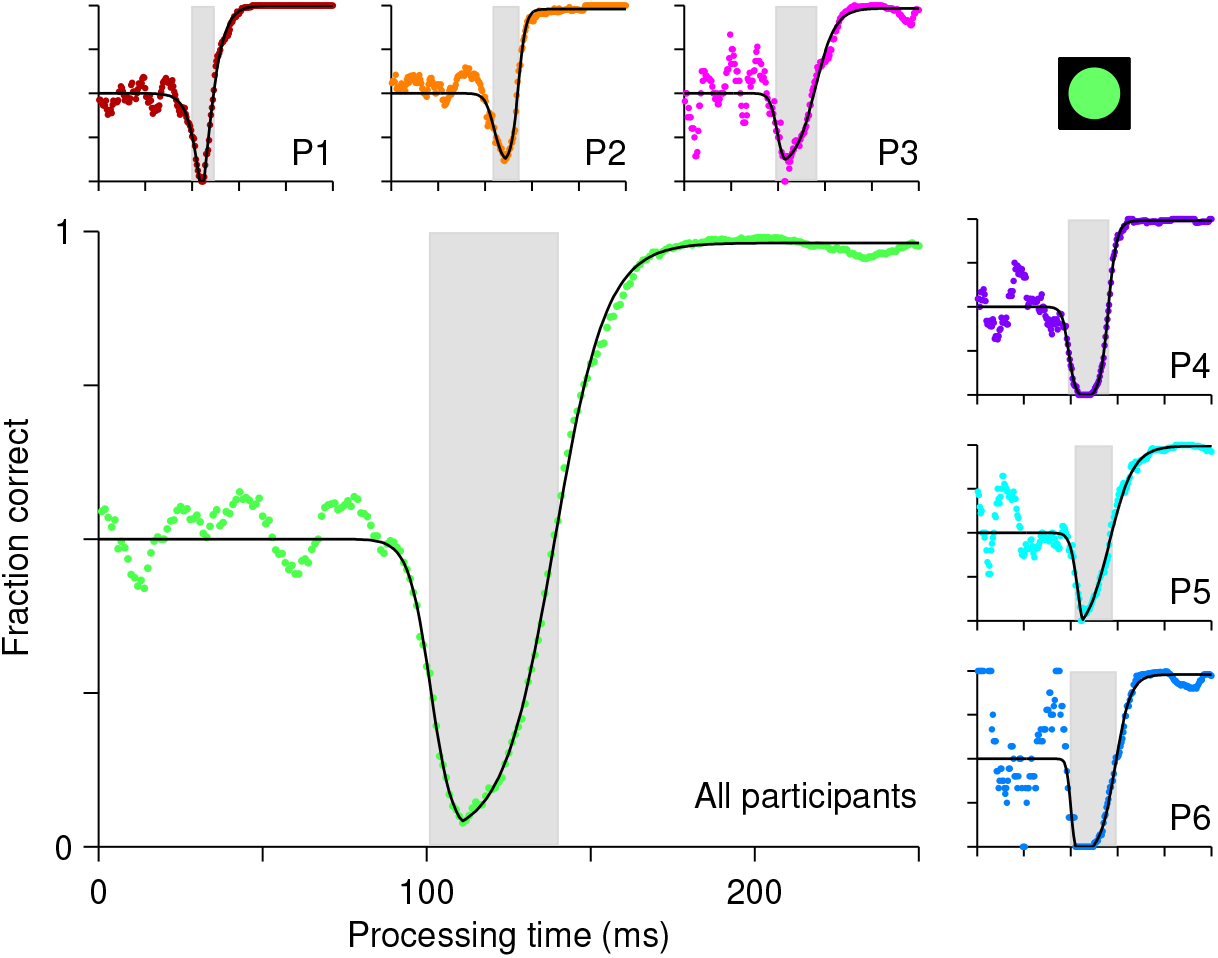
Perceptual performance in the compelled antisaccade task demonstrates an attentional vortex. Each panel shows a tachometric curve, i.e., a plot of the probability of making a correct response as a function of rPT, or cue-viewing time. Colored points are experimental results in overlapping time bins (bin width = 15 ms); black lines are continuous, analytical functions fitted to the data. The attentional vortex is the narrow rPT range over which performance drops below chance. It is demarcated by gray shades for individual participants (small panels, P1 – P6) and for the aggregate data set (large panel, All participants). Results are from trials (between 1366 and 1534 per participant) in which the high-luminance cue (icon) was shown.

Indeed, as in other perceptually-based urgent tasks (Becker and Jürgens, 1979; Stanford et al., 2010; Salinas and Stanford, 2013), the cue viewing time or rPT (computed as RT − gap, or RT + delay; Fig. 1) is the crucial variable here, because it specifies how much time is available for detecting and analyzing the cue in each trial. With little or no time to see the cue, the success rate cannot rise above chance, but as the viewing time increases, performance is expected to improve. Using multiple gap values (0–350 ms) ensures full coverage of the relevant rPT range. When the probability of making a correct choice is plotted as a function of rPT — a behavioral metric that we refer to as the tachometric curve — the result is a millisecond-by-millisecond readout of the evolving perceptual decision (Becker and Jürgens, 1979; Stanford et al., 2010; Shankar et al., 2011; Salinas and Stanford, 2013; Seideman et al., 2018).

For the compelled antisaccade task, the tachometric curve exhibits a unique, non-monotonic shape that reflects the interaction between early involuntary and later voluntary processes (Fig. 2). For rPTs shorter than 90 ms, participants perform at chance, as expected. Shortly thereafter, the initial influence of the cue manifests as a pronounced drop in performance, as participants erroneously direct a large proportion of their saccades toward the cue. This dip is short-lived (visible for rPTs of 100–140 ms approximately; Fig. 2, gray shades), but it is so abrupt and occurs so reliably over a consistent range of rPTs, that it reaches nearly 0% correct even in the data pooled from all six participants (Fig. 2, main panel). In trials in which the rPT falls inside this narrow interval — the attentional vortex — it is almost impossible to avoid looking at the cue.

As rPT increases beyond 140 ms, the success rate rises and gradually approaches an asymptote, as participants direct a progressively larger proportion of their saccades to the correct, anti location. This rise in performance is remarkable in that it is extremely fast: for the pooled data (Fig. 2, main panel) the tachometric curve goes from 0.25 to 0.75 in 18 ms, and from 0.10 to 0.90 in only 37 ms. For some of the participants the process is faster (Fig. 2, P1, P2, P4). The asymptotic fraction of correct responses is close to 1 (the lowest across participants was 0.978), which indicates that the participants understood the instructions and could perform the task almost perfectly — given enough time. Consistent with this, in easy, non-urgent antisaccade trials (Fig. 1b) the fraction correct was also close to 1 (median = 0.992). In the easy version of the task, however, processing times are typically so long that only a glimpse of the phenomenon can be found, if anything (Supplementary Fig. 1).

The characteristic shape of the tachometric curve likely results from the interplay between a reflexive and a voluntary mechanism, both of which depend on the cue. The vortex reflects the strength of low-level sensory representations that are driven by the cue’s salience and lead to attentional/oculomotor capture, whereas the recovery from the vortex reflects the strength of cognitive mechanisms that resist its pull, perform the necessary mental rotation of the cue location, and program an antisaccade. Next, to test this interpretation, we analyze how the proposed interaction varies with the salience of the cue.

### Antisaccade performance varies with cue luminance

If the vortex is indeed the result of strong, involuntary attentional capture, then it should become weaker when the salience of the cue is reduced (Theeuwes, 1991, 1992, 1994). To investigate this, our participants performed the compelled antisaccade task with cues of three luminance levels, high (data shown in Fig. 2), medium, and low (Methods). The three cues were the same for all participants, and were randomly interleaved during the experiment. Because the faintest cue was chosen to be slightly above the detection threshold, we expected it to yield a much shallower attentional vortex.

The expectation for the later rise in the tachometric curve was less clear. However, the steepness of the rise is likely to reflect, at least in part, two factors, the speed of the cognitive process that rotates the spatial location of the cue, and its variability. So, if weaker sensory signals are generally processed more slowly or with higher variance, then the tachometric curve should rise more gradually as luminance decreases.

The experimental results show that both the timing and depth of the vortex depend strongly on cue luminance. For the data pooled from all participants (Fig. 3a), as luminance decreases from high (bright green points) to low (dark green points), the vortex shifts to the right by about 50 ms (the minimum point shifts from rPT = 111 ± 1.3 ms [SE from bootstrap] to rPT = 162 ± 6.0 ms), suggesting that the time needed to detect the cue increases accordingly. The vortex also becomes much less deep (the minimum fraction correct goes from 0.03 ± 0.006 to 0.32 ± 0.026), consistent with the expected weakening of attentional capture. In addition, as luminance decreases, the rise in success rate becomes significantly less steep (p < 0.0001 for all differences in maximum slope between luminance conditions, from bootstrap; see Supplementary Fig. 2a). Thus, the salience of the cue has a substantial impact on the timing and strength of both the bottom-up and top-down mechanisms at work in the task.

Qualitatively similar dependencies on cue salience were observed in each participant’s data set (Fig. 3b), but reliable differences across individuals became evident when the effects were evaluated quantitatively. For any given tachometric curve, quantification was achieved by fitting the empirical data (Fig. 3, colored data points) with a continuous analytical function (Fig. 3, black traces; Equation 2) and measuring several features from the fitted curve (Methods). We present results for three such features that were particularly reliable given the size of our samples (for additional features, see Supplementary Fig. 2). The first one is the average value of the tachometric curve for rPTs between 0 and 250 ms, which we refer to as the mean perceptual accuracy (Fig. 4a). The second feature is the rPT at which the tachometric curve reaches its minimum, which we designate as the vortex location (Fig. 4b). And the third feature is the rPT at which the rising part of the tachometric curve is halfway between its minimum and maximum values, which we call the endogenous response centerpoint, or just the centerpoint of the curve, for brevity (Figs. 2, 3b, right border of gray shades; Fig. 4c). These quantities are partially related; the centerpoint, which measures how soon the participant can escape the vortex, is independent of the vortex location (partial Spearman correlation *ρ* = 0.34, p = 0.2; Methods), but is strongly anti-correlated with perceptual accuracy (*ρ* = −0.85, p = 10^−5^). Notably, the separation between the ‘best’ and the ‘worst’ participant within a given luminance condition is statistically large, particularly for the mean perceptual accuracy and the centerpoint of the curve (Fig. 4a, c; note little overlap between 95% confidence intervals for bars of same color). The observed effects of cue luminance are highly consistent across participants (Fig. 3b), but the quantitative details reveal idiosyncratic variations that distinguish one individual from another (Fig. 4a, c; see below). This is significant because, in general, cognitive tasks that produce robust differences between experimental conditions or treatments are typically unreliable with respect to differences between individuals (Borsboom et al., 2009; Hedge et al., 2017).

**Figure 3.**
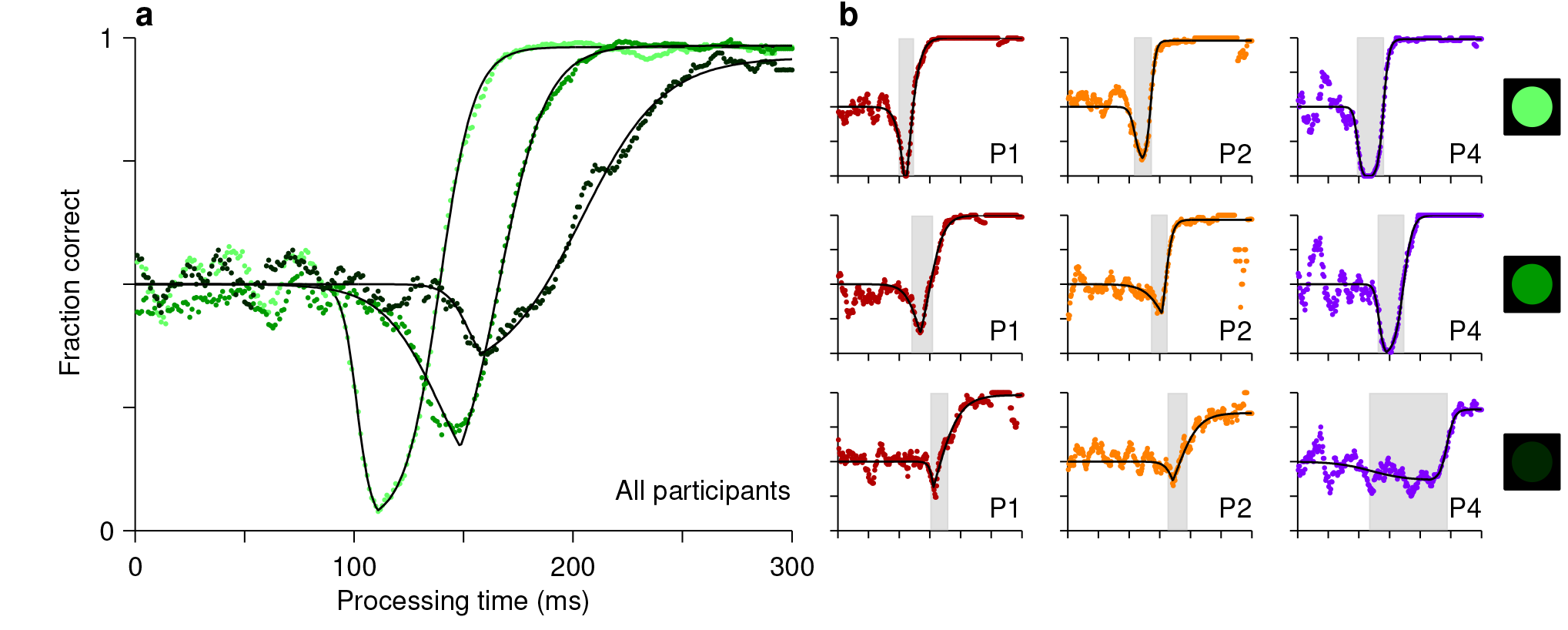
Perceptual performance varies as a function of cue luminance. **a**, Tachometric curves for trials in which the cue had high, medium, and low luminance (indicated by bright, grayish, and dark green points, respectively). Results are for the pooled data from all participants. The vortex shifts to the right and becomes less deep as luminance decreases. **b**, Tachometric curves from three individual participants at each cue luminance level, high, medium and low, as indicated by the icons. Gray shades demarcate the attentional vortex of each curve. In each panel, colored points are experimental results and black lines are continuous fits to the data.

### Individual differences in perceptual and overall performance

The tachometric curve characterizes perceptual performance in the task — i.e., how quickly and accurately the cue information can be processed. Having observed that the curve varies substantially across participants, we investigated how the individual differences in perceptual capacity relate to individual differences in more traditional antisaccade performance measures. For this, we computed the correlation between two variables (Methods), the average perceptual accuracy (mean value of the tachometric curve), and the average observed accuracy (mean fraction of correct choices). We found that, even though both quantites tend to increase with higher luminance, suggesting a positive correlation, they are, in fact, uncorrelated (Fig. 5a). The rank of a given participant based on one measure is not predictive of his or her rank based on the other.

**Figure 4.**
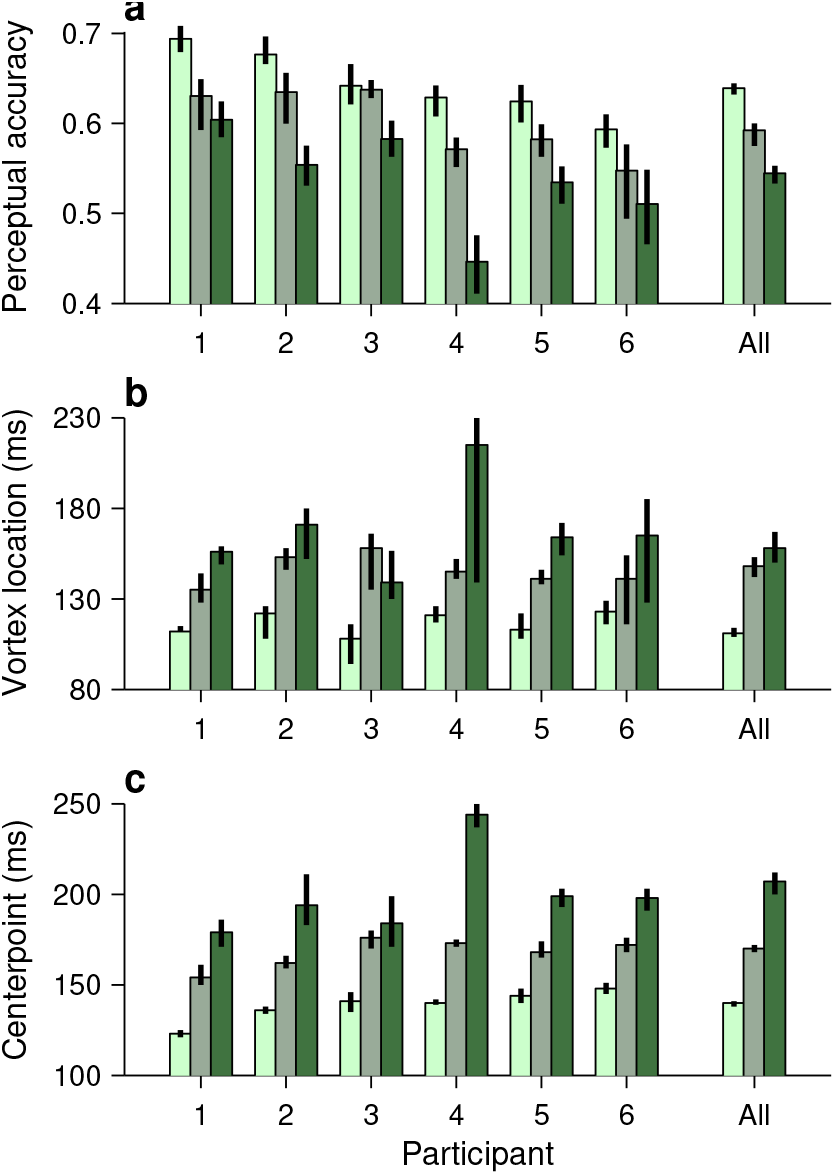
Perceptual performance quantified across participants and luminance conditions. Each panel shows one particular quantity derived from the fitted tachometric curves, with results sorted by participant (x axes) and luminance level, high (bright green), medium (grayish green), and low (dark green). Errorbars indicate 95% confidence intervals obtained by bootstrapping. **a**, Mean perceptual accuracy, calculated as the average value of the fitted tachometric curve for rPTs between 0 and 250 ms. **b**, Vortex location, calculated as the rPT at which the minimum of the fitted tachometric curve is found. **c**, Endogenous response centerpoint, equal to the rPT at which the rise of the fitted tachometric curve is halfway between its minimum and maximum values.

This may seem surprising. Logic dictates that better perceptual processing should translate into better performance — but critically, this is contingent on everything else being equal. The paradox arises because the mean RT also varies across participants, and the two accuracy measures relate to it in opposite ways. The average observed accuracy demonstrates a strong speed-accuracy tradeoff, i.e., slower participants perform better (Fig. 5b). In contrast, the mean perceptual accuracy demonstrates a weaker opposite trend; those participants that exhibit high perceptual ability also tend to respond more quickly (Fig. 5c). The results are nearly identical when perceptual performance is quantified with the endogenous response centerpoint (Supplementary Fig. 3). These comparisons show that the mean RT and accuracy — summary statistics that are commonly used to measure performance — are drastically different from the quantities that characterize perceptual or cognitive capacity in the task.

The reason for this stark divergence is that perception depends critically on cue viewing time, not on RT, whereas the overall success rate depends on both. The tachometric curve is largely independent of motor performance, as it is highly insensitive to manipulations that substantially alter the RT but not the sensory input (Stanford et al., 2010; Shankar et al., 2011; Salinas et al., 2014) (for evidence that is specific to the compelled antisaccade task, see Supplementary Fig. 4). In contrast, it is easy to see that the mean observed accuracy depends on both the shape of the tachometric curve and the subject’s urgency: because the RT and rPT are linearly related, when the RTs are predominantly short, the left side of the curve is sampled more densely and, consequently, the mean fraction correct is near chance (Fig. 2a, left of gray shade); whereas when the RTs are predominantly long, the right side of the curve is sampled more densely and the mean fraction correct is high (Fig. 2a, right of gray shade). Thus, while longer RTs have a minimal impact on perception, they cause an increase in overall success rate that is consistent with a speed-accuracy tradeoff.

**Figure 5.**
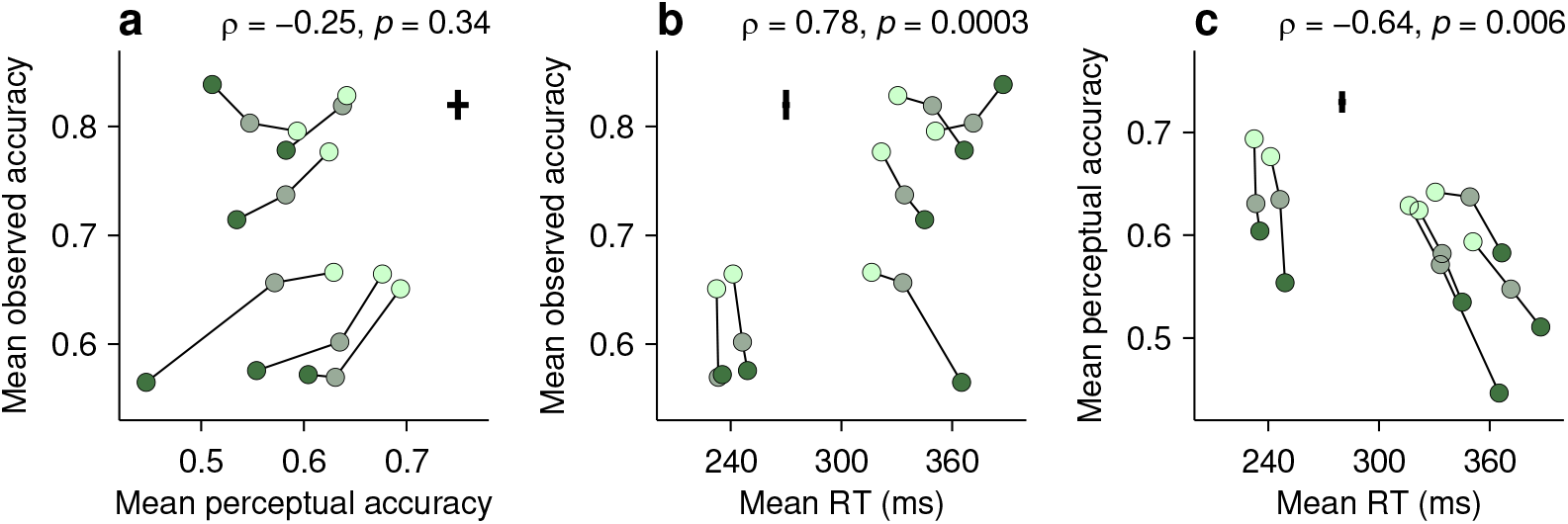
Dissociation between perceptual capacity and overall task performance. In each panel, the data from each participant (joined by lines) are shown for trials of high, medium, and low luminance cues (bright, grayish, and dark green points, respectively). Crosses indicate the typical (median) uncertainty (2 SEs) associated with the measurement in each direction. Partial Spearman correlations between values on the x and y axes are indicated, along with significance (Methods). The partial correlation eliminates the association due exclusively to luminance. **a**, Mean observed accuracy versus mean perceptual accuracy. **b**, Mean observed accuracy versus mean RT. Average RT data include both correct and incorrect trials. **c**, Mean perceptual accuracy versus mean RT.

The question of what determines the mean observed accuracy merits an additional consideration. The tachometric curve reveals three types of error: fast guesses, saccades captured by the cue, and lapses, i.e., errors found at rPT ≳ 200 ms. Lapses occur for reasons other than high motor urgency or insufficient cue viewing time, and might be indicative of distinct cognitive processes or states that vary over long time scales (Harris and Thiele, 2011; Nir et al., 2017). They were rare in the current experiment, but could conceivably be more frequent under other conditions. What should be noted is that, when the rPT is consistently extended beyond that required for asymptotic performance, as when the cue is presented before the go signal, the mean observed accuracy must primarily reflect the lapse rate (Supplementary Fig. 1). Thus, the mean observed accuracy may be informative of specific cognitive processes — just not of the ones that are normally associated with antisaccade performance.

### A comprehensive account of antisaccade behavior based on motor competition

We developed a computational model (Methods) to explore two mechanistic hypotheses about the neural origin of the vortex. This model is a variant of one that replicates both behavioral performance and choice-related neuronal activity (in the FEF) in an urgent, two-alternative, color discrimination task (Stanford et al., 2010; Shankar et al., 2011; Costello et al., 2013; Seideman et al., 2018). As in that case, the current model considers two variables, r_L_ and r_R_, that represent oculomotor responses favoring saccades toward left and right locations (Fig. 6a–c, black and red traces). These motor plans compete with each other such that the first one to reach a fixed threshold level (Fig. 6a–c, dashed lines) determines the choice: a left saccade if r_L_ reaches threshold first, or a right saccade if r_R_ reaches threshold first. In each trial, after the go signal, r_L_ and r_R_ start increasing with randomly-drawn build-up rates. The build-up process is likely to end in a random choice (i.e., a guess; Fig. 6c) when one of the initial rates is high and/or the gap is long, but otherwise, time permitting, the cue signal modifies the ongoing motor plans (Fig. 6a, b). Specifically, once the target has been identified, the plan toward it (correct) is accelerated and the other one, toward the opposite, incorrect location, is decelerated (Fig. 6a, note acceleration of black trace and deceleration of red trace after gray interval). This corresponds to the cue content, interpreted according to task rules, informing the correct choice.

**Figure 6.**
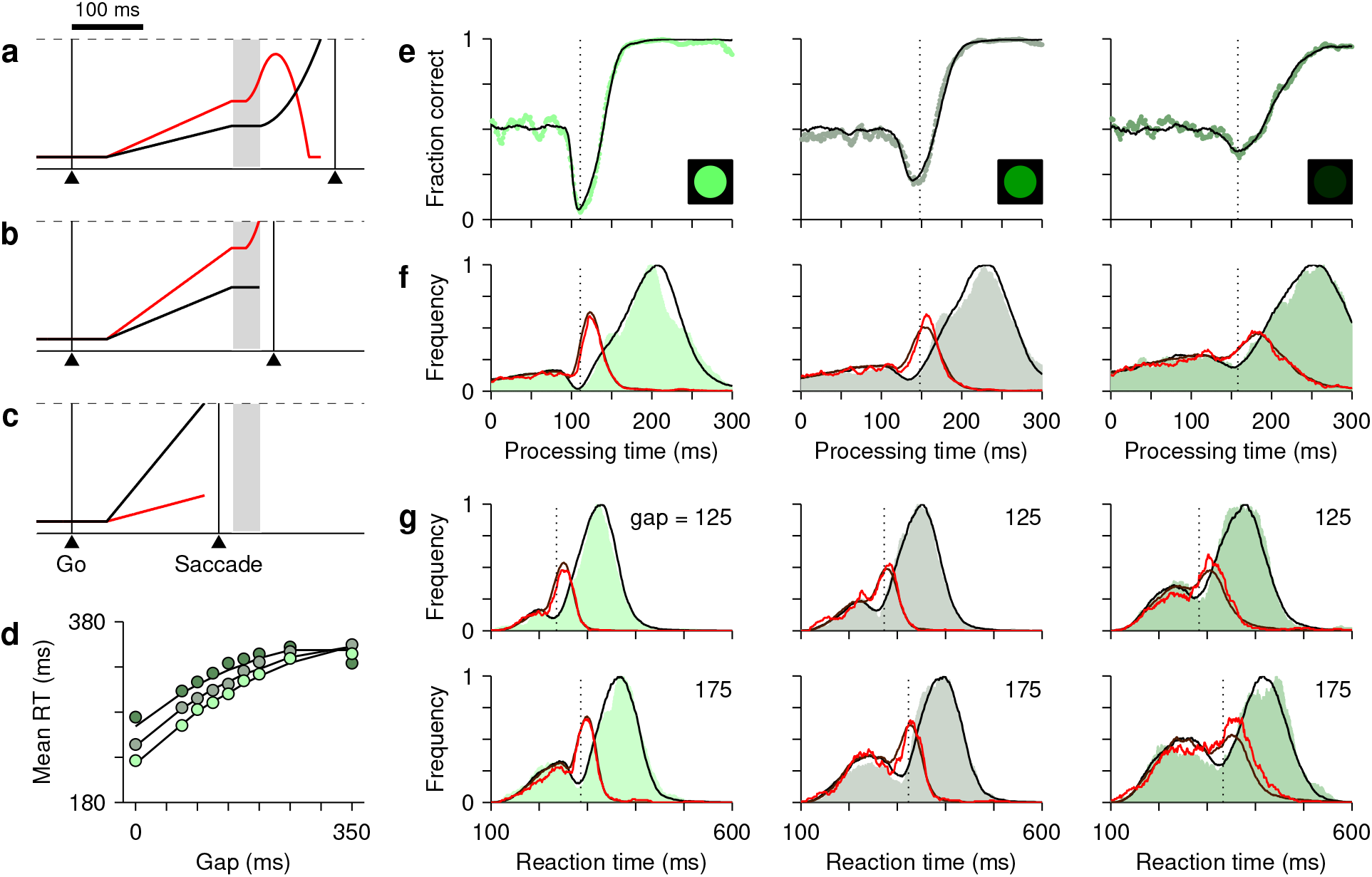
A race-to-threshold model accounts for antisaccade performance. **a**–**c**, Three simulated single trials. Traces show r_L_ and r_R_ as functions of time. These represent motor plans toward the cue (red) and toward the anti location (black). After the exogenous response interval (ERI, gray shade), the former (incorrect) plan decelerates and the latter (correct) plan accelerates. A saccade is triggered a short efferent delay after activity reaches threshold (dashed lines). In all examples, the gap is 150 ms. A correct, long-rPT trial (**a**, RT = 369 ms, rPT = 219 ms), an incorrect trial with rPT within the vortex (**b**, RT = 283 ms, rPT = 133 ms), and a correct guess (**c**, RT = 206 ms, rPT = 56 ms) are shown. The timing of the cue relative to the ongoing motor activity determines the outcome. **d**, Mean RT as a function of gap. Circles are data pooled across participants for high (bright green), medium (grayish green), and low luminance cues (dark green). SEs are smaller than symbols. Continuous lines are model results. **e**, Tachometric curves for high (left), medium (middle), and low (right) luminance cues. Continuous lines are model results. **f**, Processing time distributions for correct (shades) and incorrect trials (red traces) at each luminance level. Overlaid traces (black and dark red) are corresponding model results. **g**, RT distributions for correct (shades) and incorrect trials (red traces) at individual gap values (125 and 175 ms) for each luminance level. Overlaid traces (black and dark red) are model results. Dotted vertical lines in **e**–**g** mark the rPT at which the vortex reaches its minimum point (vortex location). All empirical data are pooled across participants.

To adapt this “accelerated race-to-threshold” model to the compelled antisaccade task, we introduced one crucial, task-specific assumption: that the competition is biased in favor of the cue location during a period of time that we refer to as the exogenous response interval, or **ERI** (Fig. 6a–c, gray shades). During the ERI, the cue has already been detected by the circuit but not yet interpreted as “opposite to the target” (so it cannot yet drive the acceleration and deceleration described above). We consider two possible mechanisms by which, during the ERI, the detection of the cue may lead to exogenous attentional/oculomotor capture: (1) it could halt or suppress the ongoing plan toward the anti location (Fig. 6a, b, black traces during gray interval), or (2) it could transiently accelerate the ongoing plan toward the cue location (Fig. 6a, b, red traces during gray interval). These alternatives are not mutually exclusive. The former is consistent with evidence that salient, abrupt-onset stimuli reflexively interrupt ongoing saccade plans (Dorris et al., 2007; Bompas and Sumner, 2011; Hafed and Ignashchenkova, 2013; Buonocore et al., 2017; Salinas and Stanford, 2018), whereas the latter is consistent with the short-latency, stimulus-driven activation of visually responsive neurons in oculomotor areas (Gottlieb and Goldberg, 1999; Thompson et al., 2005; Ipata et al., 2006; Joiner et al., 2017; White et al., 2017; Chen et al., 2018).

We found that, to reproduce the psychophysical data accurately, both mechanisms were necessary. To see why, first note that the tachometric curve, which refers to the proportion of correct choices in each rPT bin, can be expressed as a ratio, 
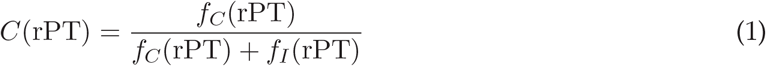
where *f_C_*(rPT) and *f_I_*(rPT) describe the frequencies of correct and incorrect choices at each rPT, i.e., they are the rPT distributions for correct and incorrect trials (normalized by the same factor; Methods). Each of these distributions demonstrates a distinct feature: *f_C_* (Fig. 6f, green shades) has a dip, whereas *f_I_* (Fig. 6f, red traces) has a peak. Both features contribute to the attentional vortex, as dictated by the above expression. The critical mechanistic observation is that acceleration of the motor plan toward the cue during the ERI accounts for the peak in *f_I_*, whereas interruption of the competing motor plan away from the cue produces the dip in f_C_. Thus, when the model was implemented with either one of the mechanisms alone, it failed to replicate the experimental feature associated with the other (Supplementary Fig. 5). However, with the two mechanisms acting simultaneously, in coordination, the model reproduced the full data set in quantitative detail.

First, for the data pooled across participants, the model fitted the tachometric curve (Fig. 6e) and the rPT distributions for correct and incorrect responses (Fig. 6f). Second, for individual gap conditions, the model reproduced not only the variations in mean success rate and RT (Fig. 6d), but also the shapes of the RT distributions for correct and incorrect choices, which were typically bimodal (Fig. 6g). Third, the model accurately captured all the dependencies on luminance (Fig. 6e–g, compare results across columns). Importantly, in doing so, the values of the parameters that correspond to pure motor performance (the distribution of initial build-up rates for r_L_ and r_R_, and the distribution of afferent delays associated with the go signal) were the same across luminance conditions (Table 1), in correspondence with the fact that all trials proceeded identically up to cue presentation, and that trials with different gap and cue luminance were interleaved during the experiment. And fourth, the model also fitted the (noisier) data from individual participants, even though they showed large, idiosyncratic variations in motor performance, as well as in the dips and peaks of their rPT distributions (Supplementary Figs. 6, 7). In terms of mean RT and accuracy, and of the characteristic features of the tachometric curve, the empirical (Figs. 4, 5) and simulated data sets (Supplementary Figs. 8, 9) were nearly indistinguishable.

Mechanistically, the best-fitting parameter values of the model (Table 1) provide further insight about the crucial element that gives rise to the attentional vortex — the exogenous bias during the ERI (Fig. 6a–c, gray shades). Consider the following values based on the fits to the pooled data. According to the model, the onset of the ERI, which corresponds to the time at which the cue is detected, is highly sensitive to luminance. For the high, medium, and low conditions, the oculomotor circuitry detects the cue 76 ± 5 ms (mean ± SD for simulated trials), 104 ± 13 ms, and 126 ± 19 ms after its presentation. These detection times determine the vortex location, and their difference from high to low luminance (50 ms) corresponds closely with the rightward shift of the vortex observed experimentally (51 ms; Fig. 3a). Remarkably, the ERI lasts only 24 ms (on average) in all three conditions, and the exogenous acceleration of the plan toward the cue occurs only during the last 14 ms (high luminance), or only during the last 10 ms (medium and low luminance); before that, the plan toward the cue halts just like its counterpart toward the anti location (Fig. 6a, b, note that red trace is initially flat during gray interval). The model suggests that the exogenous acceleration favoring the cue location is very brief but very powerful, which explains why the left edge of the vortex can be so steep.

Finally, the parameter values (Supplementary Table 1) also point to specific neural mechanisms that likely underlie the individual differences in perceptual capacity. In general, identifying those mechanisms is complicated because their variations across (random) participants and across (controllable) experimental conditions are not necessarily correlated (Borsboom et al., 2009; Hedge et al., 2017). The cue latency discussed above is a perfect example: it demonstrates (via parameter 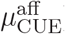) a strong, consistent dependence on luminance in each participant’s data set, and yet, for a given luminance level, it is not predictive of individual perceptual accuracy (Supplementary Fig. 10a). By contrast, we hypothesize that the magnitudes of the exogenous and endogenous acceleration (via parameters a_EX_ and a_END_) are major sources of individual variation, because although they have weaker dependencies on luminance, they are reliable predictors of perceptual accuracy (Supplementary Fig. 10b–d).

## Discussion

The current work is significant because it achieves a marked improvement in the measurement of fundamental psychophysical quantities. By design, the antisaccade task creates a conflict between exogenous and endogenous mechanisms, the former driven by the saliency of the cue and the latter by task instructions followed willfully. Other tasks (e.g., Kim and Cave, 1999), most notably the singleton-distracter task employed by Theeuwes and colleagues (Theeuwes, 1991, 1992, 1994; Theeuwes et al., 1998, 1999; Nissens et al., 2017), have been highly successful at revealing such conflict in the form of attentional or oculomotor capture, but they require more complex visual displays with multiple items and a secondary discrimination to serve as a probe of the effect, all leading to RTs that are much longer (≫ 250 ms) than typical intersaccadic intervals. In principle, a minimalistic task typifying such an essential phenomenon would be extremely useful; it could serve to determine the neural correlates of volitional versus reflexive action, or pinpoint the consequences of disease states on specific cognitive abilities, for example. Numerous studies based on the traditional antisaccade task have, in fact, reported large differences in overall performance between distinct populations of participants (Guitton et al., 1985; Klein and Foerster, 2001; Munoz et al., 2003; Condy et al., 2007; Hakvoort et al., 2012; Antoniades et al., 2015) — but without relating them directly to the conflict at the heart of the task. It is now clear why: the interaction is so fast, that the standard behavioral metrics, mean accuracy and RT, are blind to it. By making the task urgent, focusing on processing time (instead of RT), and developing a mechanistic model that is firmly grounded on the neurophysiology of saccadic choices, we were able to resolve the opposing influences of endogenous and exogenous mechanisms on the oculomotor response with unprecedented sharpness.

The results make four main contributions. First, they identify concrete ways in which endogenous and exogenous mechanisms act on the oculomotor circuitry — namely, via acceleration, deceleration, and halting of rising firing rates — together with their unique behavioral signatures (dips and peaks in rPT distributions). Second, they characterize the time scales of those mechanisms (a few tens of ms) as well as their dependencies on luminance. Third, they clearly parse the contributions of motor (RT) and perceptual performance (tachometric curve) to the probability of success in each trial, thus removing the pervasive confounds caused by the speed-accuracy tradeoff. And fourth, they relate variations in motor, perceptual, and cognitive mechanisms to individual differences in performance. In particular, all participants exhibited a vortex, but different vortices resulted from different combinations of exogenous and endogenous mechanisms (compare rPT distributions for P1 vs. P4 in Supplementary Figs. 6, 7) — a degeneracy in neural function that is reminiscent of that found in more reduced circuits (Marder et al., 2015).

### The attentional vortex as a perceptuo-motor phenomenon

The shape of the main behavioral metric in the task, the tachometric curve, can be qualitatively understood by realizing that the rPT conveys information not only about cue viewing time, but also about the initial build-up rates in each trial (as depicted in Fig. 6a–c). Short-rPT trials (< 90 ms with high luminance) occur when at least one of the initial motor plans rises quickly, so a saccade is typically triggered before the cue information has had time to reach the circuit and impact the outcome (Fig. 6c). Conversely, long-rPT trials (> 150 ms with high luminance) result when the initial motor activity builds-up slowly, in which case the exogenous bias is not sufficient to push it above threshold, and the choice is guided by the later, fully-processed cue information (Fig. 6a). In the intermediate case (100 ≲ rPT ≲ 140 ms), however, rPTs fall in a Goldilocks region in which, by the time the cue is detected, the initial motor activity toward the cue is still below threshold but already so high, that the exogenous bias typically propels it past threshold before the endogenous modulation can act (Fig. 6b).

Previous studies support key elements of this account. First, single-neuron recordings in monkeys show that the initial level of motor activity contributes as proposed: during urgent saccadic choices, the build-up of activity in FEF starts after the go signal, regardless of when the cue information arrives (Stanford et al., 2010; Costello et al., 2013), and during antisaccade performance, the ongoing activity in FEF and SC is higher before erroneous saccades toward the cue than before correct antisaccades (Everling et al., 1998; Everling and Munoz, 2000). Second, the timing of the vortex and its dependence on luminance parallel those of cue-driven visual bursts in the oculomotor system (Gottlieb and Goldberg, 1999; Thompson et al., 2005; Ipata et al., 2006; Joiner et al., 2017; White et al., 2017; Chen et al., 2018). And third, the sudden presentation of a salient distracter stimulus has a robust impact on a developing saccade plan, with the effect depending strongly on their spatial congruence: when the saccade target is diametrically opposite to the distracter, there is ample evidence (reviewed by Salinas and Stanford, 2018) indicating that the developing plan is transiently halted or suppressed, whereas when the saccade target is near the abrupt-onset stimulus, the developing plan is boosted (Dorris et al., 2007; Edelman and Xu, 2009; White et al., 2013; Marino et al., 2015). These observations are consistent with the two exogenous biasing mechanisms implemented by the model and, more generally, with the notion that the attentional vortex arises from a particular combination of external (cue exposure) and internal (motor urgency) conditions that is highly conducive to oculomotor capture.

### Coupling between spatial attention and saccade planning

Our results are pertinent to a mechanistic question that is central to the “premotor theory” of attention: to what degree is the neural substrate of the deployment of spatial attention the same as that of saccade planning? There is strong evidence that planning a saccade to a given point in space automatically implies that attentional resources are at least partially allocated to that point (Kowler et al., 1995; Moore and Fallah, 2001; Godijn and Theeuwes, 2003; Cavanaugh and Wurtz, 2004; Steinmetz and Moore, 2014; Klapetek et al., 2016). However, the converse relationship — i.e., whether spatial attention can be deployed without necessarily planning a saccade — has been more contentious. Some studies suggest that the answer is “yes” (Juan et al., 2004; Thompson et al., 2005). Others, however, indicate that the hypothesized motor plan associated with attentional allocation is just very difficult to observe when fixation must be actively maintained (Belopolsky and Theeuwes, 2012). During fixation, such a plan may manifest only as a subtle increase in baseline neural activity, rather than via the more typical steady rise in firing rate (Hauser et al., 2018), but it can be uncovered through experimental manipulations (Theeuwes et al., 1998, 1999; Katnani and Gandhi, 2013; Nissens et al., 2017), and is evident in microsaccades (Chen et al., 2015; Lowet et al., 2018). Our results are consistent with the idea that the deployment of exogenous spatial attention is equivalent to a reflexive increase in saccade-related activity (in, say, the FEF). When a salient cue is detected, a bias favoring a motor plan toward its location is always generated (with the bias consisting of acceleration of the plan toward the cue and halting of any plans away from it). However, the impact of the exogenous biasing signal is limited, because it is very brief and can generally be counteracted by endogenous mechanisms — except when the oculomotor circuitry is in just the right state, i.e., when activity related to the next saccade is still below threshold and directionally ambiguous, but has already developed to a substantial degree. Then the exogenous bias becomes observable as an overt, captured saccade.

## Methods

### Subjects and setup

Experimental subjects were six healthy human volunteers, two male and four female, ages 21– 30. All had normal or corrected-to-normal vision. All participants provided informed written consent before the experiment. All experimental procedures were conducted with the approval of the Institutional Review Board (IRB) of Wake Forest School of Medicine.

The experiments took place in a semi-dark room. The participants sat on an adjustable chair, with their chin and forehead supported, facing a VIEWPixx LED monitor (VPixx Technologies Inc., Saint Bruno, Quebec, Canada; 1920 × 1200 screen resolution, 120 Hz refresh rate, 12 bit color) at a distance of 52 cm. Viewing was binocular. Eye position was recorded using an EyeLink 1000 infrared camera and tracking system (SR Research, Ottawa, Canada) with a sampling rate of 1000 Hz. Stimulus presentation was controlled using the system’s integrated software package (Experiment Builder).

### Behavioral tasks

The sequence of events in the antisaccade task is described in Fig. 1. The inter-trial interval was 1 s. The gap values used were −200, −100, 0, 75, 100, 125, 150, 175, 200, 250, and 350 ms, where negative numbers correspond to delays in the easy antisaccade task (Fig. 1b). Thus, compelled and easy, non-urgent trials were interleaved. In each trial, the gap value, cue location (−10^°^ or 10^°^), and luminance level (see below) were randomly sampled. Auditory feedback was provided at the end of each trial: a beep to indicate that the saccadic response was made within the allowed RT window (450 ms), or no sound if the limit was exceeded. This feedback was independent of the choice. The task was run in blocks of 150 trials. After 50–150 trials of practice, each participant completed 30 blocks over 6 experimental sessions (days). Within each session, 2–3 minutes of rest were allowed between blocks.

The cue was a green circle (0.5^°^ diameter) appearing on a black background. Each participant performed the task with cues of three luminance levels, high (17.6 cd m^−2^), medium (0.35 cdm^−2^), and low (0.22 cd m^−2^) Luminance was measured with a spectrophotometer (i1 Pro 2 from χ-Rite, Inc., Grand Rapids, MI). The cues were generated in Adobe Illustrator using the 8-bit RGB vectors [15 168 40], [3 28 7], and [1 12 3]. The lowest luminance was chosen to be close to the detection threshold based on detection curves generated previously for two participants.

### Data analysis

All data analyses were carried out using customized scripts written in Matlab (The MathWorks, Natick, MA). Except where explicitly noted, results are based on the analysis of urgent trials (gap ≥ 0) only; i.e., easy trials (delay trials with gap < 0) were excluded.

In each trial, the rPT was computed by subtracting the gap value from the RT value recorded in that trial. We refer to this processing time as ‘raw’ because it includes any afferent or efferent delays in the circuitry (Stanford et al., 2010). To compute the tachometric curve and rPT distributions, trials were grouped into rPT bins of 15 ms, with bins shifting every 1 ms. Normalized rPT distributions, *f_C_*(rPT) and *f_I_*(rPT), were obtained by counting the numbers of correct and incorrect trials, respectively, in each rPT bin, and dividing both functions by the same factor. The tachometric curve, which gives the proportion of correct trials in each bin, was then computed using Equation 1. For display purposes, the normalization factor used was the maximum value of *f_C_* or *f_I_*, whichever was largest, but the factor has no effect on the tachometric curve.

In order to quantify perceptual performance, each tachometric curve was fitted with a continuous analytical function, *v*(*x*), which was defined as 
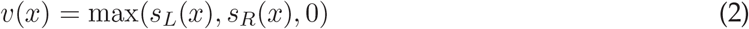
where themaximum function max(*a, b, c*) returns *a, b*, or *c*, whichever is largest, and *s_L_* and *s_R_* are two sigmoidal curves. These are given by 
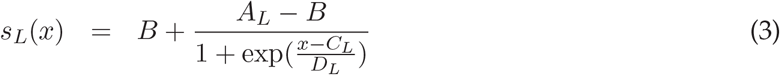

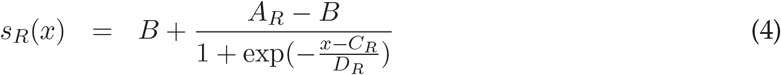
where *s_L_* tracks the left (decreasing) side of the tachometric curve and *s_R_* tracks the right (increasing) side. The asymptotic value on the left side was fixed at *A_L_* = 0.5, to enforce the constraint that, for very short processing times, performance must be at chance. For any given empirical tachometric curve, the six remaining parameters defining *v*, the coefficients *B, A_R_, C_L_, C_R_, D_L_*, and *D_R_*, were adjusted to minimize the mean absolute error between the experimental and fitting functions. The minimization was done using the Matlab function `fminsearch`.

Once the best-fitting *v*(rPT) function for a given tachometric curve was found, we numerically calculated eight quantities, or features, from it: the asymptotic value (equal to *A_R_*), the minimum value (vortex depth), the rPT at which the minimum was found (vortex location), the most negative slope, the most positive slope, the rPT for which *v* is exactly between 0.5 (chance) and the minimum (left edge of the curve), the rPT for which *v* is exactly between the minimum and the asymptote (the curve’s centerpoint), and the average of the curve for rPTs between 0 and 250 ms (mean perceptual accuracy). In Figs. 2, 3, the gray shades demarcating the vortex correspond to the interval between the left edge and the centerpoint of each tachometric curve. Confidence intervals for all of these quantities were obtained by bootstrapping (Davison and Hinkley, 2006; Hesterberg, 2014); that is, by resampling the data with replacement and recalculating all the quantities many times to generate distributions for them. This was done in five steps: (1) resample the original trials with replacement, keeping the original number of contributing trials, (2) recompute the empirical tachometric curve from the resampled trials, (3) fit the new tachometric curve with a continuous *v* function, (4) recompute the eight characteristic features from the new *v* function, and (5) repeat steps 1–4 10,000 times to generate distributions for all the features. Reported confidence intervals correspond to the 2.5 and 97.5 percentiles obtained from the bootstrapped distributions.

To quantify the association between average quantities computed for individual participants, such as the mean RT or mean observed accuracy (Fig. 5), we considered three data points per participant, one for each luminance condition. The strength of association and its significance were calculated with three methods. First we computed the partial Pearson correlation coefficient, which is the standard linear correlation between two variables but controlling for the effect of a third one, luminance in this case. This was implemented via the Matlab function `partialcorr`. We also computed the partial Spearman correlation coefficient, which involves a similar calculation but based on the ranks of the data points. This was using `partialcorr too`. Finally, using the Matlab function `fitlm`, we fitted the data to a linear regression model that also included luminance as a variable. The three methods typically produced similar results. We report those obtained with the partial Spearman correlation, which is denoted as *ρ*, because they were generally the most conservative.

### The accelerated race-to-threshold model

The model for the compelled antisaccade task is a straightforward extension of one developed previously for a two-alternative, urgent, color discrimination task (Salinas and Stanford, 2010; Shankar et al., 2011; Costello et al., 2013; Seideman et al., 2018). As explained in the main text, the idea is that two motor plans (in the FEF), represented by firing rate variables *r_L_* and *r_R_*, compete with each other to trigger a saccade toward a left or a right location. To describe the model and its parameters, it is useful to re-label these rate variables as *r_C_* and *r_A_*, where the subscripts now refer to the cue and anti locations (keeping in mind that the *C* and *A* labels are randomly assigned to left and right directions in each trial). Over time, these plans advance toward a fixed threshold (equal to 1000 arbitrary units, or **AU**). If *r_C_* exceeds threshold first, the saccade is incorrect, toward the cue, whereas if *r_A_* exceeds threshold first, the saccade is correct, away from the cue. The saccade is considered to be triggered a short efferent delay (equal to 20 ms) after threshold crossing.

The two rate variables evolve as follows 
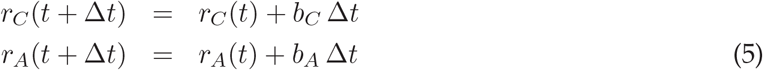
where *b_C_* and *b_A_* are their respective build-up rates and the time step Δ*t* is equal to 1 ms. When the build-up rates are constant, the firing rates r_C_ and r_A_ increase linearly over time. Periods during which the activity accelerates or decelerates are those during which the build-up rates themselves change steadily, as described below. Any negative *r_C_* and *r_A_* values are reset to zero. Each simulated trial can be subdivided into three epochs with different model dynamics.

<u>Epoch 1</u>: before the ERI. Each trial starts with the two activity variables, *r_C_* and *r_A_*, equal to zero. The go signal occurs at *t* = 0, but the two motor plans start building up later, after an afferent delay. This afferent delay is drawn from a Gaussian distribution with mean 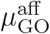 and SD 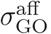, where values below 20 ms are excluded. The initial build-up rates, 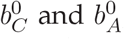, are drawn from a two-dimensional Gaussian distribution with mean *μ_b_*, SD *σ_b_*, and correlation coefficient *ρb*. During this epoch, after the initial afferent delay has elapsed, *r_C_* and *r_A_* evolve according to Equations 5, with 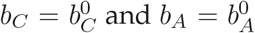. If during this period one of the motor plans exceeds the threshold, a saccade is produced and the trial ends. Otherwise, the trial continues.

**Table 1.** Parameters of the race-to-threshold model for the pooled data. Build-up rates are in AU ms^−1^ times are in ms, and acceleration and deceleration are in AU ms^−2^

| lum | $\mu_b$ | $\sigma_b$ | $\rho_b$ | $\mu_{GO}^{aff}$ | $\sigma_{GO}^{aff}$ | $\mu_{CUE}^{aff}$ | $\sigma_{CUE}^{aff}$ | $\mu_{ERI}$ | $\sigma_{ERI}$ | $g_{ERI}$ | $\Delta_{ERI}$ | $a_{EX}$ | $d_{END}$ | $a_{END}$ | $\lambda$ |
| --- | --- | --- | --- | --- | --- | --- | --- | --- | --- | --- | --- | --- | --- | --- | --- |
| high | 1.4 | 3.74 | -0.95 | 51 | 36 | 76 | 5 | 24 | 4 | 0 | 10 | 0.96 | -0.70 | 0.17 | 0.02 |
| med | 1.4 | 3.74 | -0.95 | 51 | 36 | 104 | 13 | 24 | 3 | 0 | 14 | 1.15 | -0.54 | 0.17 | 0.02 |
| low | 1.4 | 3.74 | -0.95 | 51 | 36 | 126 | 19 | 24 | 10 | 0 | 14 | 0.58 | -0.29 | 0.14 | 0.10 |

<u>Epoch 2</u>: during the ERI. The start of the ERI corresponds to the time point at which the cue is detected by the model circuit (we stress that this is a local event, and make no claims about the participant’s perceptual experience). Cue detection occurs after an afferent delay relative to the time of cue presentation, which is at *t* = gap. This delay is drawn from a Gaussian distribution with mean 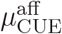 and 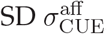, where values below 20 ms are excluded. The duration of the ERI also varies normally across trials. It is drawn from a Gaussian distribution with mean μ_ERI_ and SD *σ_ERI_*, with negative values reset to zero. The two motor plans behave differently during the ERI. For the plan toward the anti location, *r_A_*, the build-up rate of is 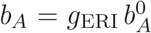, where the constant gain factor *g_ERI_* is either zero (i.e., the plan halts) or negative (i.e., the plan is suppressed). This factor was set to zero for the pooled data, but negative values were allowed when fitting the data from individual participants. Whether zero or negative, the build-up rate of the anti plan is the same throughout the whole ERI. In contrast, for the motor plan toward the cue, *r_C_*, the build-up rate is 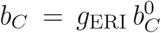 but only during the first Δ_ERI_ ms of the ERI; thereafter this build-up rate instantly recovers its initial value (so 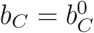) and then increases steadily, such that 
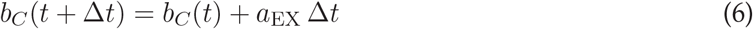
until the end of the ERI, where the term *a_EX_* is the exogenous acceleration of the cue plan. In this way, the plan toward the cue, *r_C_*, first halts for Δ_ERI_ ms and then accelerates. If *r_C_* exceeds threshold during the ERI, a saccade toward the cue is triggered. Otherwise, the trial continues.

<u>Epoch 3</u>: after the ERI. During this last period, the plan toward the anti location first recovers its initial value (instantly, so 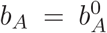) and then accelerates, whereas the plan toward the cue decelerates. That is, 
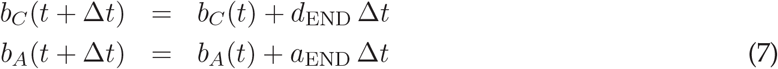
where the endogenous deceleration *d_END_* is negative and the endogenous acceleration *a_END_* is positive. The process continues until one of the plans reaches the threshold.

Finally, the model also considers lapses, trials in which errors are made for reasons other than insufficient cue viewing time. Lapses occur with a probability λ, and are implemented as trials in which the endogenous acceleration and deceleration are equal to zero. In other words, a lapse corresponds to a trial in which the information about the correct target never reaches the circuit. During lapses, after the ERI (epoch 3), the motor plan toward the anti location continues building up at its initial rate, 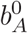, whereas the plan toward the cue continues advancing at whatever buildup rate it achieved at the end of the ERI.

In all, the model has 15 parameters that were adjusted to fit the pooled data set or the data from individual participants. Best-fitting values are listed in Table 1 and Supplementary Table 1. These were obtained by searching over a multidimensional parameter space, gradually reducing its volume, seeking to minimize the mean absolute error between the simulated and the experimental data. For each parameter vector tested, the error consisted of a sum of terms, each representing one target function to be fitted. These functions were the RT distributions for correct choices at individual gaps, the RT distributions for incorrect choices, also at individual gaps, and the tachometric curve. The search/minimization procedure was repeated multiple times with different initial conditions to ensure that solutions were found near the global optimum.

A Matlab script for running the model is provided as supplementary material.

---

**Supplementary Figure 1.**
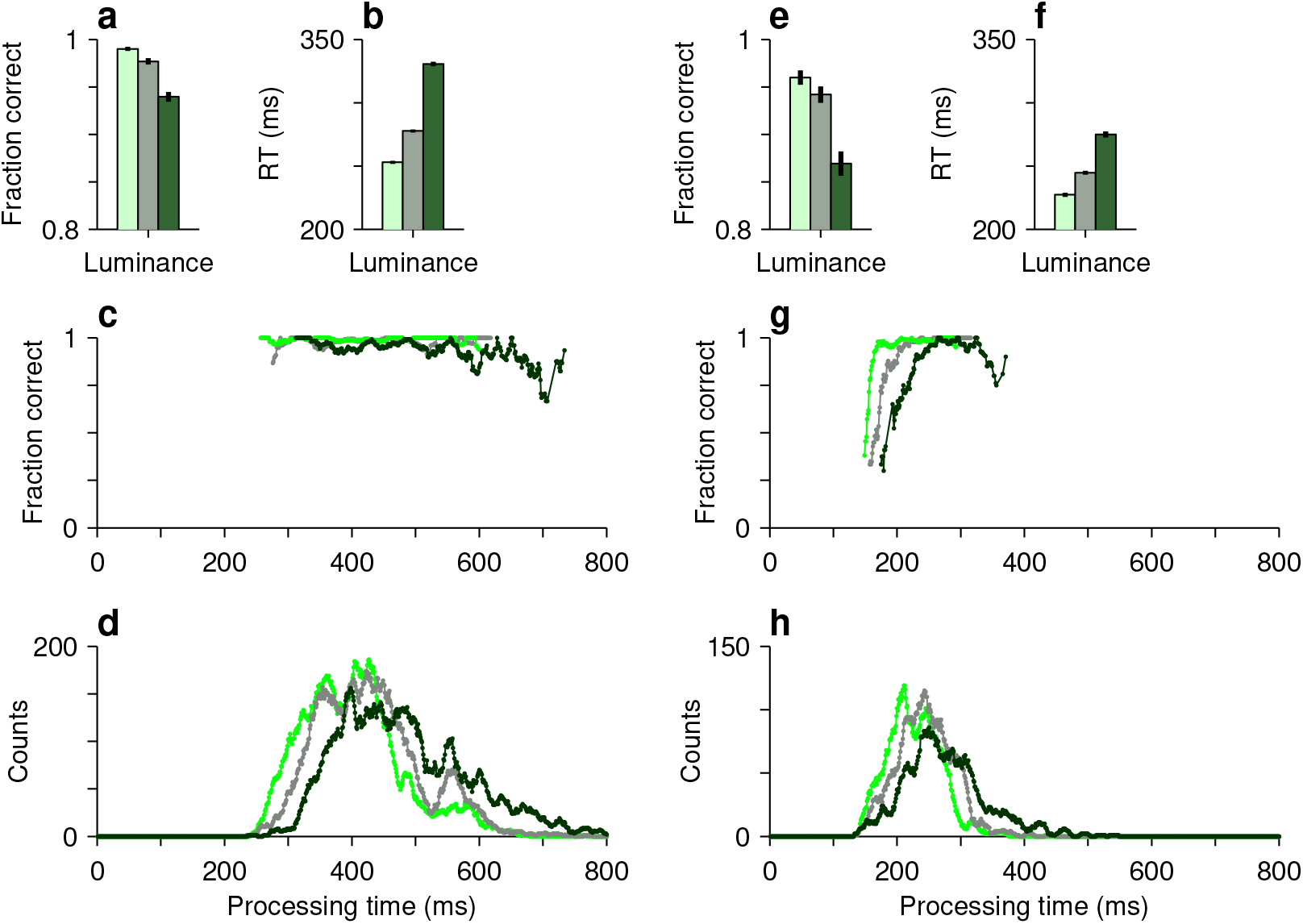
Performance in non-urgent antisaccade trials. All data are pooled across participants and sorted by luminance level, high (bright green), medium (grayish green), and low (dark green). Results are shown separately for easy trials (left side), in which the delay between cue and go onset was 100 or 200 ms, and zero-gap trials (right side), in which cue and go onset were simultaneous. **a**, Mean fraction correct in easy trials (± 1 SE, from binomial proportion). **b**, Mean RT in easy trials (± 1 SE across trials). **c**, Tachometric curves from easy trials. **d**, Processing time distributions in easy trials. Both correct and incorrect responses are included. **e**–**h**, As in **a**–**d**, except for zero-gap trials. In easy trials, essentially all rPTs correspond to asymptotic performance. In zero-gap trials, the late rise in the tachometric curve becomes partially observable, but the majority of rPTs still correspond to asymptotic performance.

**Supplementary Figure 2.**
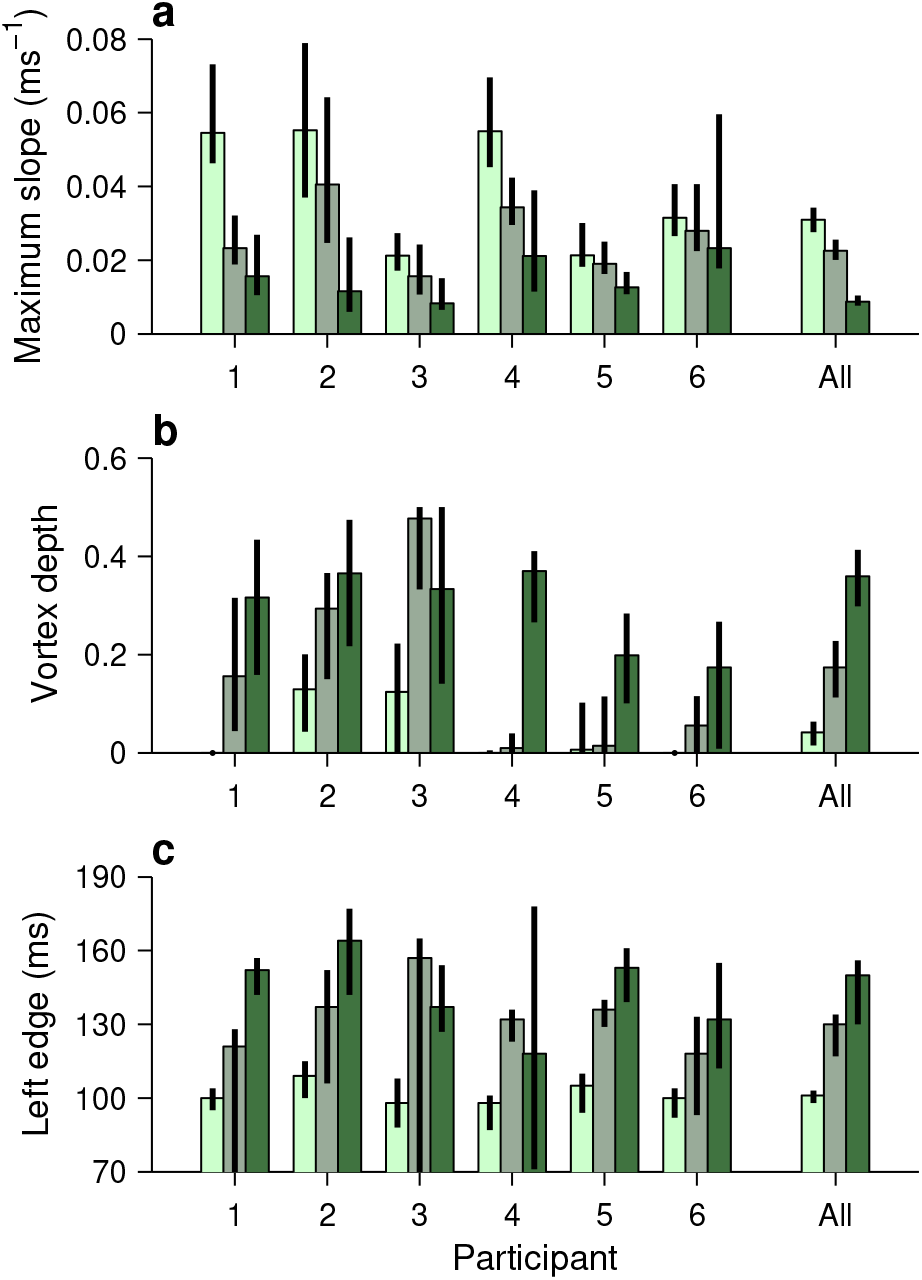
Additional quantities that characterize perceptual performance across participants and cue conditions. Each panel shows one particular feature derived from the fitted tachometric curves, with results sorted by participant (x axes) and luminance level, high (bright green), medium (grayish green), and low (dark green). Errorbars indicate 95% confidence intervals obtained by bootstrapping. Same format as in Fig. 4. **a**, Maximum slope attained during the rising phase of the tachometric curve. **b**, Vortex depth, equal to the minimum fraction correct. Note that, for participants 1, 4, and 6, the depth is zero for the high-luminance cue. **c**, Left edge of the vortex, equal to the rPT at which the dip of the tachometric curve is halfway between chance (0.5) and its minimum value. Although these metrics are somewhat noisier than those shown in Fig. 4, the effects of luminance are again highly consistent across participants, albeit with significant quantitative differences.

**Supplementary Figure 3.**
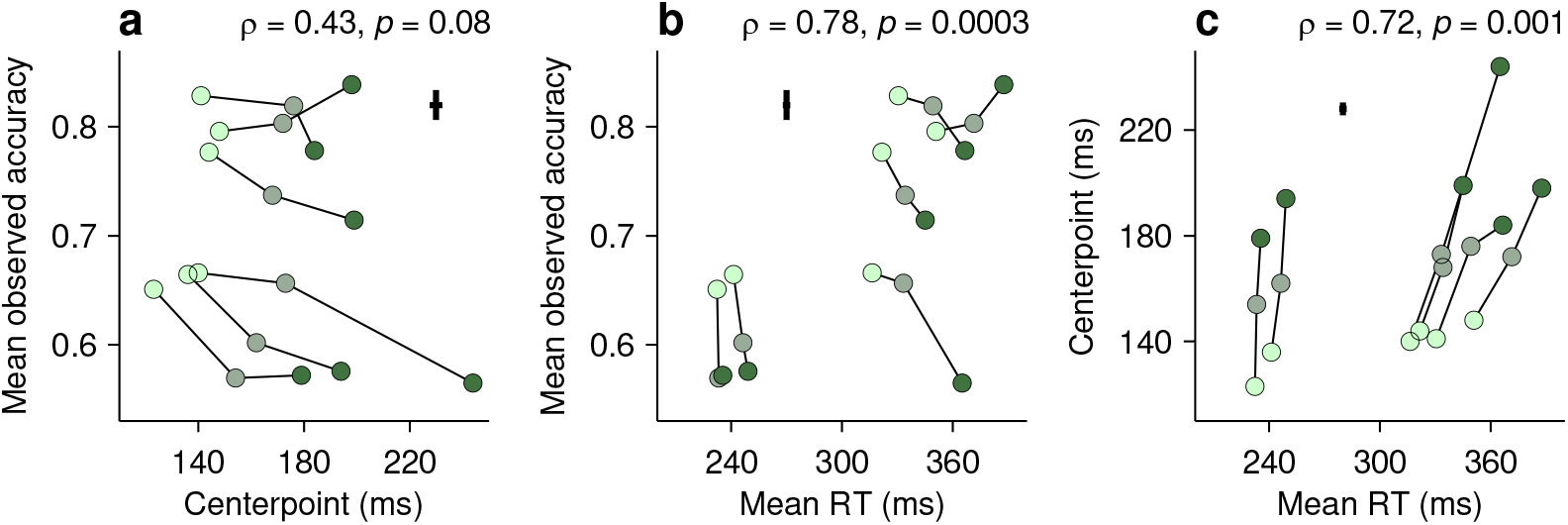
Dissociation between perceptual capacity and overall task performance. Same format as in Fig. 5, except that the endogenous response centerpoint is plotted instead of the mean perceptual accuracy. **a**, Mean observed accuracy as a function of the centerpoint. **b**, Mean observed accuracy as a function of mean RT. **c**, Centerpoint as a function of mean RT. The centerpoint relates to the mean observed accuracy and the mean RT (this figure) in nearly identical ways as the mean perceptual accuracy (Fig. 5), if one takes into account that a *lower* centerpoint corresponds to better perceptual performance.

**Supplementary Figure 4.**
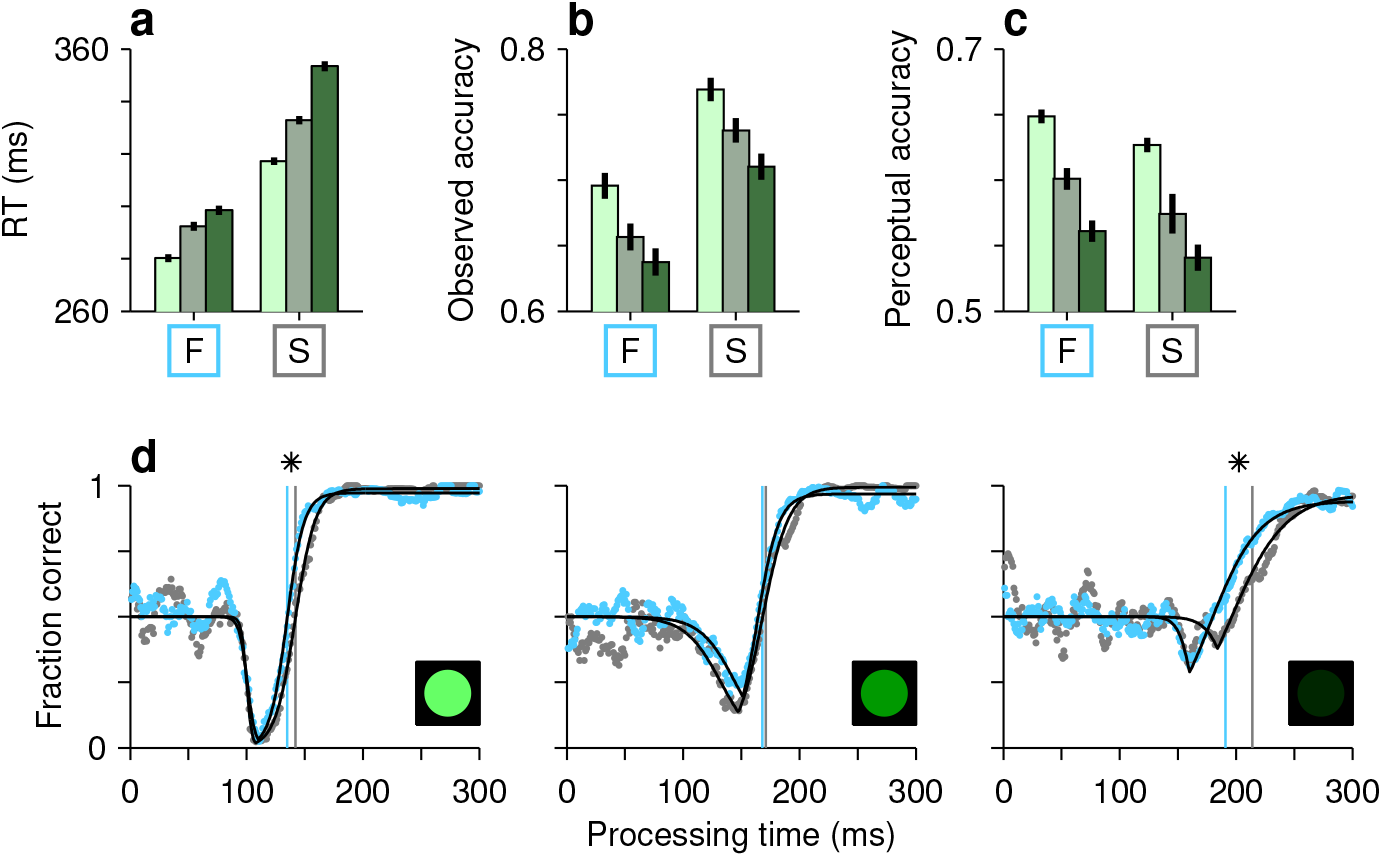
Decoupling perceptual and motor performance. All results are based on data aggregated across participants. Each panel compares results in fast (F) versus slow (S) blocks of trials, which were sorted as follows. First, the 30 blocks of trials performed by each participant were ranked, post hoc, according to mean RT. Then the 10 blocks at the top and the 10 at the bottom were designated as the fast and slow blocks, respectively. Finally, the fast and slow blocks were separately pooled across participants to yield two corresponding groups of trials. **a**, Mean RT (± 1 SE) in compelled antisaccade trials. Bar colors, bright, grayish, and dark green, correspond to cue luminance, high, medium, and low, respectively. **b**, Mean observed accuracy, equal to the measured fraction of correct responses (± 1 SE from binomial proportion), in compelled antisaccade trials. Note that, for a given luminance, the observed accuracy was higher in the slow than in the fast blocks. **c**, Mean perceptual accuracy, equal to the mean value of the fitted tachometric curve over the 0–250 ms rPT range (± 1 SE from bootstrap). Note that perceptual accuracy was slightly lower in the slow than in the fast blocks. **d**, Tachometr
ic curves for fast (cyan dots) and slow blocks (gray dots). Each panel shows data for a specific cue luminance level, as indicated by the icons. Vertical lines mark the centerpoints of the fitted curves. In the high- and low-luminance cases, the blue curve is slightly shifted to the left relative to the gray (asterisks indicate p < 0.0005 for the difference in centerpoints, from bootstrap). This implies slightly better perceptual performance in the fast than in the slow blocks.

**Supplementary Figure 5.**
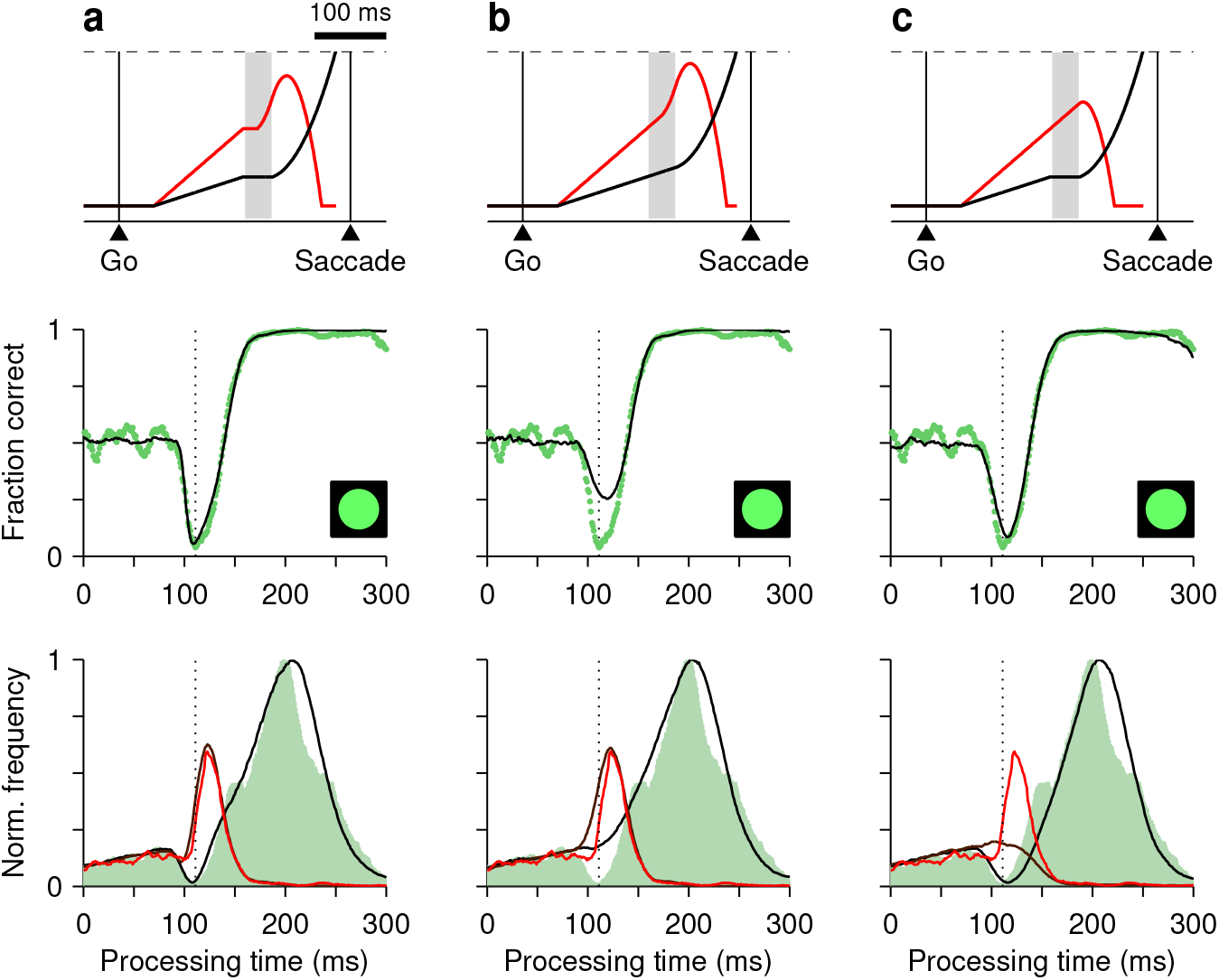
Contributions of two distinct neural mechanisms to attentional/oculomotor capture. Results are for three versions of the race-to-threshold model. For each one, a single-trial example (top row), the tachometric curve (middle row), and the rPT distributions (bottom row) are shown. All data are from the high luminance condition. For each model, the simulation results were obtained with the parameter values that minimized the error between the model and the pooled experimental data (Methods). **a**, Results from the full model in which, during the ERI (gray shade, top panel), the motor plan toward the anti location (black trace, top panel) halts and that toward the cue (red trace, top panel) halts briefly and then accelerates. The empirical tachometric curve (green dots, middle panel) is shown together with the simulated curve (black trace, middle panel). The empirical rPT distributions for correct (green shade, bottom panel) and incorrect responses (red trace, bottom panel) are shown together with the simulated distributions (black and dark red traces, bottom panel). Same data as in Fig. 6e, f, left column. The full model reproduces both rPT distributions. **b**, As in **a**, but for a restricted version of the model in which, during the ERI, the motor plan toward the cue accelerates but the plan toward the anti location keeps building up at its initial rate. The transient cue acceleration alone generates an appropriate peak in the rPT distribution for errors, but cannot reproduce the dip in the rPT distribution for correct choices. As a consequence, the simulated vortex is not deep enough. **c**, As in **a**, but for another restricted version of the model in which, during the ERI, the motor plan toward the anti location halts but that toward the cue keeps advancing, unperturbed. By itself, the transient interruption of the plan away from the cue accounts for the dip in the rPT distribution for correct choices and is sufficient to produce a vortex with appropriate depth. However, it cannot replicate the prominent peak in the rPT distribution for incorrect choices.

**Supplementary Figure 6.**
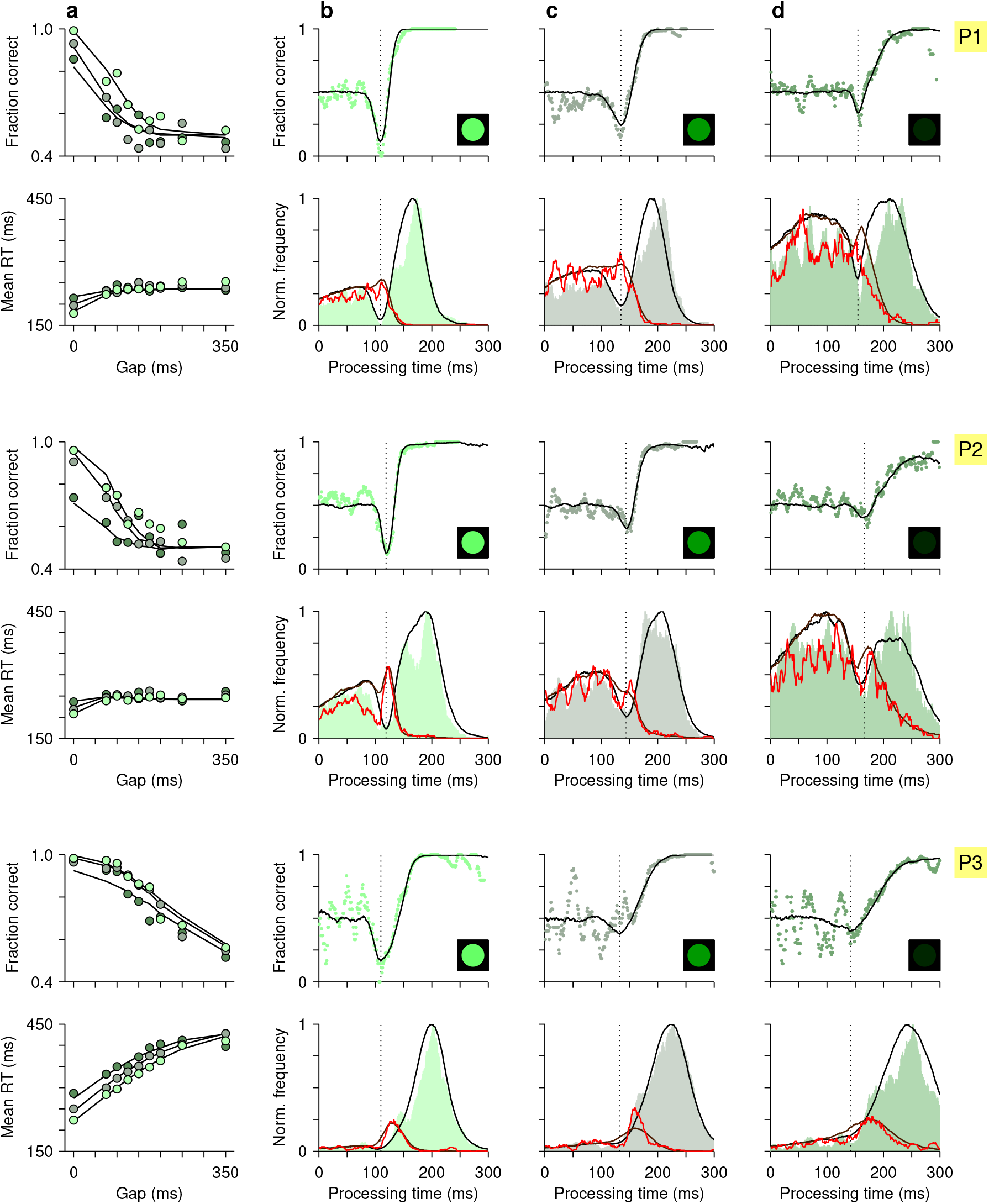
The accelerated race-to-threshold model individually fitted to the data of participants 1 (top two rows), 2 (middle rows), and 3 (bottom two rows), as indicated on the right. **a**, Mean fraction of correct choices (top) and mean RT (bottom) as functions of gap. Circles are experimental results for high (bright green), medium (grayish green), and low luminance (dark green) conditions. Continuous lines are model results. **b**–**d**, Tachometric curves for high (**b**), medium (**c**), and low luminance cues (**d**) are shown in the top panels. Corresponding rPT distributions for correct (shades) and incorrect trials (red traces) are shown in the bottom panels. Black and dark-red continuous lines are model results. For each luminance, the dotted vertical line marks the vortex location.

**Supplementary Figure 7.**
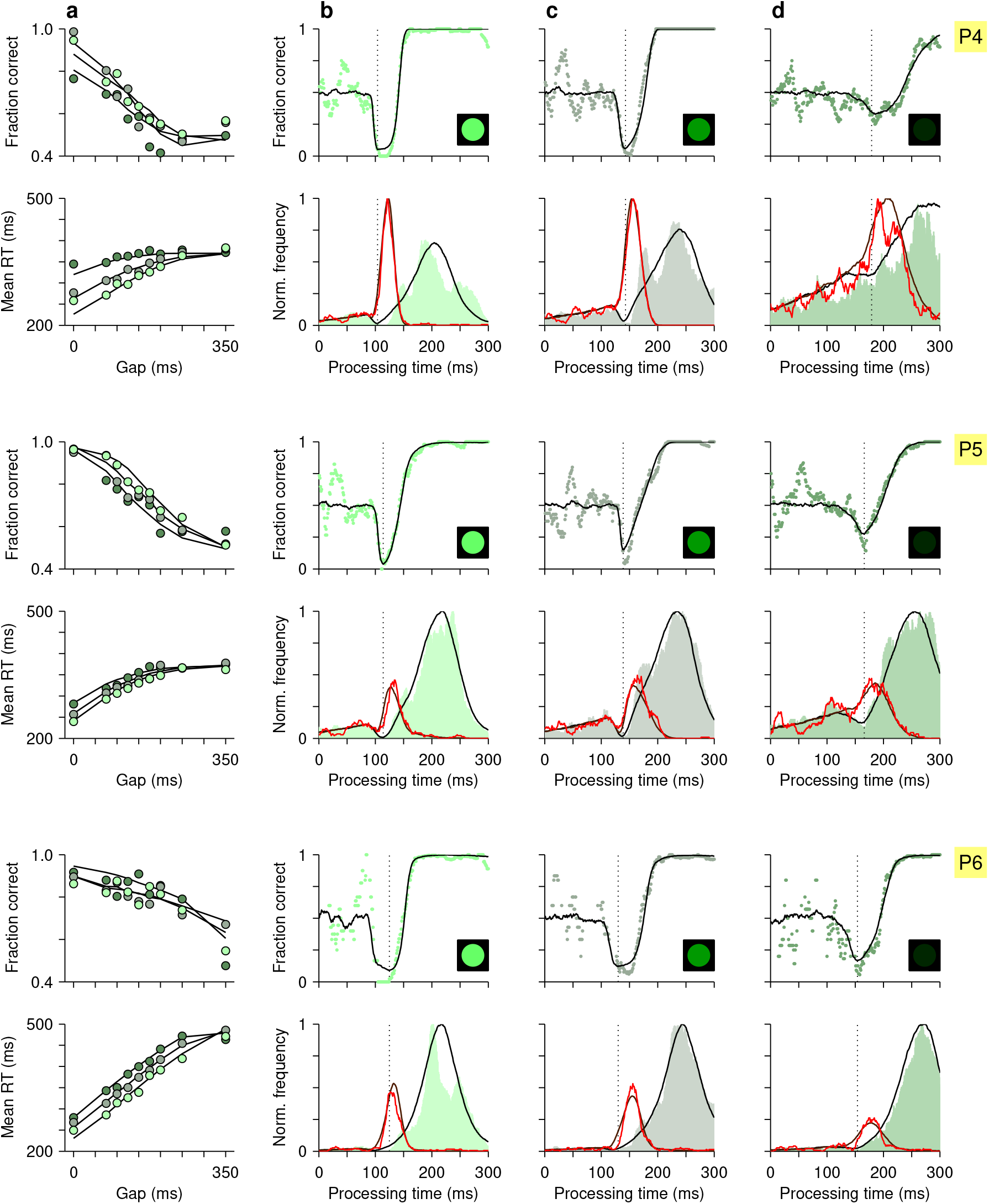
The accelerated race-to-threshold model individually fitted to the data of participants 4 (top two rows), 5 (middle rows), and 6 (bottom two rows). Same format as in Supplementary Fig. 6.

**Supplementary Figure 8.**
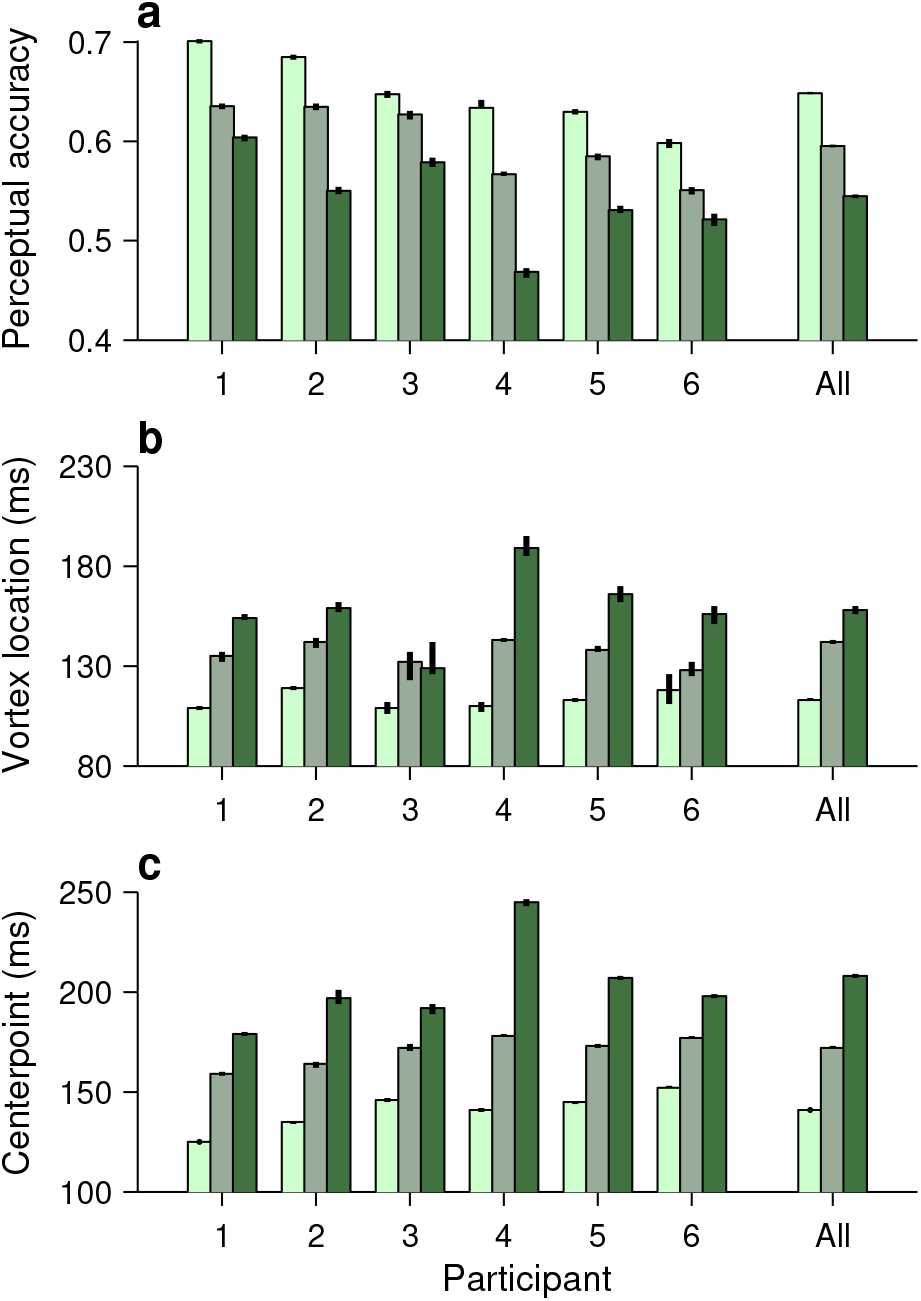
Simulated perceptual performance quantified in the same way as the experimental data. Results were obtained with the exact same analyses as those in Fig. 4 (Methods), except that they were based on simulated trials. The model data were generated using the best-fitting model parameters for each individual participant. Same format as in Fig. 4.

**Supplementary Figure 9.**
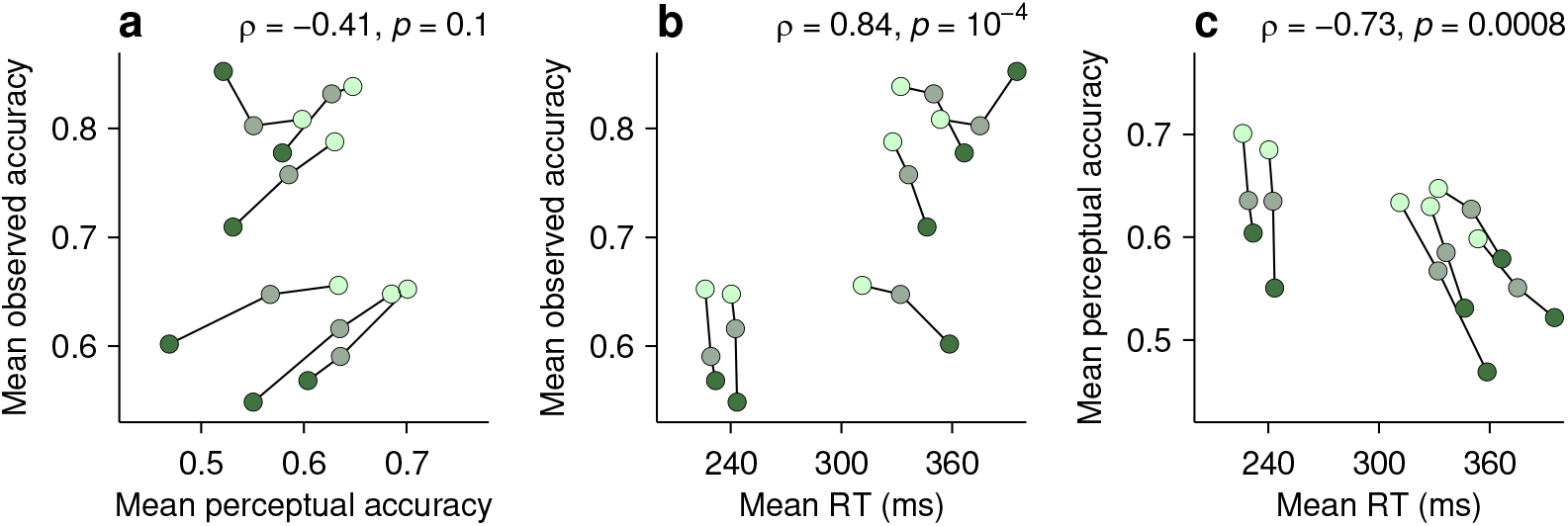
Dissociation between perceptual capacity and overall task performance as seen with the accelerated race-to-threshold model. Results were obtained in the exact same way as those in Fig. 5, except that they were based on simulated trials. The model data were generated using the best-fitting model parameters for each individual participant. Same format as in Fig. 5.

**Supplementary Figure 10.**
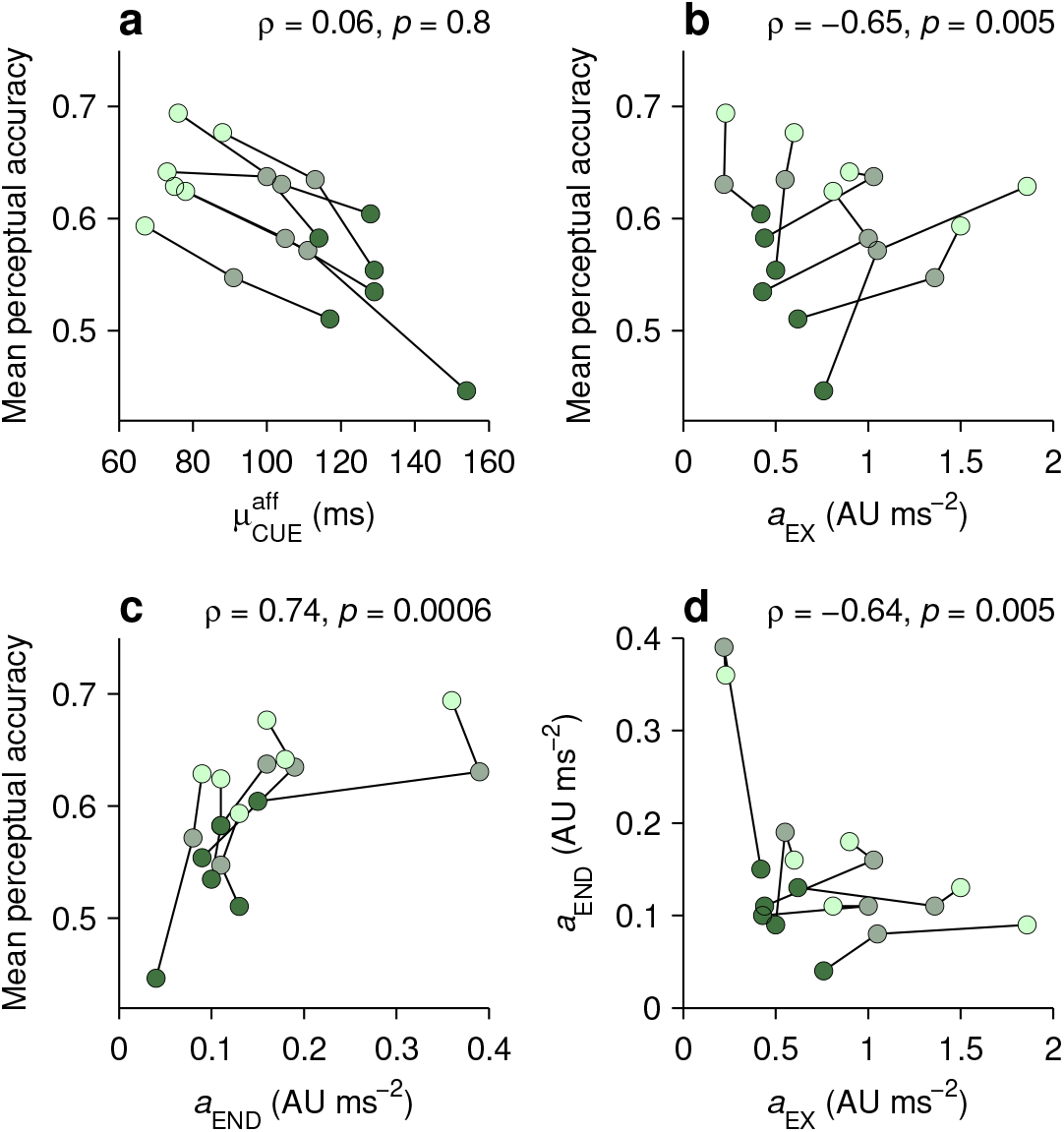
Model parameters that characterize individual perceptual performance. In each panel, the data from each participant (joined by lines) are shown for trials of high, medium, and low luminance cues (bright, grayish, and dark green points, respectively). Parameter values are as in Supplementary Table 1. Partial Spearman correlations between values on the x and y axes are indicated, along with significance (Methods). The partial correlation eliminates the association due to luminance. **a**, Mean perceptual accuracy as a function of the mean afferent delay of the cue (parameter 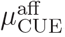, or mean cue latency). There seems to be a strong, negative association, but it is all due to mutual dependencies on luminance; within each luminance level, cue latency does not predict perceptual accuracy. This is an example of Simpson’s paradox, a statistical effect whereby a trend between two variables disappears or reverses when the data are conditioned on a third variable. **b**, Mean perceptual accuracy as a function of the exogenous acceleration (parameter *a*_EX_). **c**, Mean perceptual accuracy as a function of the endogenous acceleration (parameter *a*_END_). **d**, Correlation between endogenous and exogenous acceleration. The negative relationship indicates that participants whose attention is captured strongly by the cue also tend to endogenously shift their visuospatial attention more slowly. Other model parameters showed weaker relationships to perceptual accuracy.

**Supplementary Table 1.** Parameters of the race-to-thresholdmodel for individual participants. Participant (P1–P6) and luminance (high, medium, low) are indicated in the first column. For comparison, the results for the aggregate data (A) are shown at the top.

| P/lum | $\mu_b$ | $\sigma_b$ | $\rho_b$ | $\mu_{GO}^{aff}$ | $\sigma_{GO}^{aff}$ | $\mu_{CUE}^{aff}$ | $\sigma_{CUE}^{aff}$ | $\mu_{ERI}$ | $\sigma_{ERI}$ | $g_{ERI}$ | $\Delta_{ERI}$ | $a_{EX}$ | $d_{END}$ | $a_{END}$ | $\lambda$ |
| --- | --- | --- | --- | --- | --- | --- | --- | --- | --- | --- | --- | --- | --- | --- | --- |
| A/high | 1.4 | 3.74 | -0.95 | 51 | 36 | 76 | 5 | 24 | 4 | 0 | 10 | 0.96 | -0.70 | 0.17 | 0.02 |
| A/med | 1.4 | 3.74 | -0.95 | 51 | 36 | 104 | 13 | 24 | 3 | 0 | 14 | 1.15 | -0.54 | 0.17 | 0.02 |
| A/low | 1.4 | 3.74 | -0.95 | 51 | 36 | 126 | 19 | 24 | 10 | 0 | 14 | 0.58 | -0.29 | 0.14 | 0.10 |
| P1/high | 4.1 | 4 | -0.92 | 48 | 40 | 76 | 8 | 20 | 2 | -0.6 | 2 | 0.36 | -0.60 | 0.23 | 0.00 |
| P1/med | 4.1 | 4 | -0.92 | 48 | 40 | 104 | 15 | 24 | 2 | -0.5 | 1 | 0.39 | -0.58 | 0.22 | 0.00 |
| P1/low | 4.1 | 4 | -0.92 | 48 | 40 | 128 | 4 | 9 | 8 | -0.1 | 3 | 0.15 | -0.20 | 0.42 | 0.02 |
| P2/high | 4.5 | 2.37 | -0.81 | 44 | 34 | 88 | 7 | 16 | 1 | -0.5 | 6 | 0.60 | -0.90 | 0.16 | 0.03 |
| P2/med | 4.5 | 2.37 | -0.81 | 44 | 34 | 113 | 13 | 17 | 2 | -0.8 | 8 | 0.55 | -0.80 | 0.19 | 0.05 |
| P2/low | 4.5 | 2.37 | -0.81 | 44 | 34 | 129 | 10 | 11 | 13 | -0.4 | 7 | 0.50 | -0.64 | 0.09 | 0.36 |
| P3/high | 1.7 | 1.7 | -0.91 | 47 | 40 | 73 | 11 | 26 | 6 | 0 | 15 | 0.90 | -0.59 | 0.18 | 0.01 |
| P3/med | 1.7 | 1.7 | -0.91 | 47 | 40 | 100 | 20 | 22 | 2 | 0 | 17 | 1.03 | -0.18 | 0.16 | 0.00 |
| P3/low | 1.7 | 1.7 | -0.91 | 47 | 40 | 114 | 29 | 22 | 9 | 0 | 10 | 0.44 | -0.27 | 0.11 | 0.07 |
| P4/high | 2.6 | 1.34 | -0.97 | 55 | 41 | 75 | 2 | 14 | 8 | -0.9 | 5 | 1.86 | -1.05 | 0.09 | 0.00 |
| P4/med | 2.6 | 1.34 | -0.97 | 55 | 41 | 111 | 4 | 17 | 6 | -0.1 | 4 | 1.05 | -0.72 | 0.08 | 0.00 |
| P4/low | 2.6 | 1.34 | -0.97 | 55 | 41 | 154 | 22 | 16 | 6 | 0.0 | 9 | 0.76 | -0.22 | 0.04 | 0.15 |
| P5/high | 2.3 | 1.82 | -0.92 | 54 | 35 | 78 | 6 | 24 | 5 | -0.4 | 12 | 0.81 | -0.68 | 0.11 | 0.03 |
| P5/med | 2.3 | 1.82 | -0.92 | 54 | 35 | 105 | 5 | 21 | 4 | -0.2 | 16 | 1.00 | -0.22 | 0.11 | 0.00 |
| P5/low | 2.3 | 1.82 | -0.92 | 54 | 35 | 129 | 19 | 24 | 7 | -0.4 | 11 | 0.43 | -0.23 | 0.10 | 0.00 |
| P6/high | 0.5 | 1.9 | -0.95 | 46 | 47 | 67 | 8 | 20 | 14 | -0.6 | 13 | 1.50 | -1.27 | 0.13 | 0.03 |
| P6/med | 0.5 | 1.9 | -0.95 | 46 | 47 | 91 | 7 | 17 | 9 | -0.3 | 9 | 1.36 | -0.65 | 0.11 | 0.04 |
| P6/low | 0.5 | 1.9 | -0.95 | 46 | 47 | 117 | 12 | 24 | 12 | -0.1 | 7 | 0.62 | -0.87 | 0.13 | 0.02 |

Author contributions
ES and TRS designed the research; BRS, LS, SMF, DDA, and CKH collected and analyzed data with guidance from ES and TRS; ES performed modeling work; ES and TRS wrote the manuscript.

## Abbreviations

AU: arbitrary units
ERI: exogenous response interval
FEF: frontal eye field
rPT: raw processing time
RT: reaction time
SC: superior colliculus
SD: standard deviation
SE: standard error

